# German-wide interlaboratory study compares consistency, accuracy and reproducibility of whole-genome short read sequencing

**DOI:** 10.1101/2020.04.22.054759

**Authors:** Laura Uelze, Maria Borowiak, Erik Brinks, Carlus Deneke, Kerstin Stingl, Sylvia Kleta, Simon H. Tausch, Kathrin Szabo, Anne Wöhlke, Burkhard Malorny

**Affiliations:** Department of Biological Safety, German Federal Institute for Risk Assessment (BfR), Max-Dohrn-Str. 8-10, 10589 Berlin, Germany; Department of Microbiology and Biotechnology, Max Rubner-Institut (MRI), Herrmann-Weigmann-Str. 1, 24103 Kiel, Germany; Department 5, Federal Office of Consumer Protection and Food Safety (BVL), Diedersdorfer Weg 1, 12277 Berlin, Germany; Food and Veterinary Institute, Lower Saxony State Office for Consumer Protection and Food Safety (LAVES), Dresdenstr. 2, 38124 Braunschweig, Germany

**Author notes:** Corresponding Author, German Federal Institute for Risk Assessment, Study Centre for Genome Sequencing and Analysis, Department Biological Safety, Diedersdorfer Weg 1, 12277 Berlin, Germany.

**Keywords:** interlaboratory study, whole-genome sequencing, food safety

## Abstract

We compared the consistency, accuracy and reproducibility of next-generation short read sequencing between ten laboratories involved in food safety (research institutes, state laboratories, universities and companies) from Germany and Austria. Participants were asked to sequence six DNA samples of three bacterial species (*Campylobacter jejuni, Listeria monocytogenes* and *Salmonella enterica*) in duplicate, according to their routine in-house sequencing protocol. Four different types of Illumina sequencing platforms (MiSeq, NextSeq, iSeq, NovaSeq) and one Ion Torrent sequencing instrument (S5) were involved in the study. Sequence quality parameters were determined for all data sets and centrally compared between laboratories. SNP / and cgMLST calling were performed to assess the reproducibility of sequence data collected for individual samples. Overall, we found Illumina short read data to be more accurate and consistent than Ion Torrent sequence data, with little variation between the different Illumina instruments. Two laboratories with Illumina instruments submitted sequence data with lower quality, probably due to the use of a library preparation kit, which shows difficulty in sequencing low GC genome regions. Differences in data quality were more evident after assembling short reads into genome assemblies, with Ion Torrent assemblies featuring a great number of allele differences to Illumina assemblies. Clonality of samples was confirmed through SNP calling, which proved to be a more suitable method for an integrated data analysis of Illumina and Ion Torrent data sets, than cgMLST calling.

## 1. Introduction

Whole genome sequencing (WGS) is a high resolution, high-throughput method for the molecular typing of bacteria. Through bioinformatic analysis of bacterial genome sequences, it is not only possible to identify bacteria on a species and sub-species level, but also to identify antimicrobial resistance and virulence genes. Further, it is possible through a variety of methods, such as variant calling, k-mer based, or gene-by-gene approaches, to determine the relatedness / clonality between bacterial isolates, making it the ideal tool for outbreak studies, routine surveillance and clinical diagnostics (Ronholm et al., 2016). Initially expensive and difficult to set up, the technology is becoming continuously more user-friendly and affordable (Uelze et al., 2020). In recent years, funding provided through federal initiatives has enabled public health and food safety laboratories in Germany and worldwide to acquire sequencing platforms. A number of different sequencing technologies exist, each with their own upsides and shortcomings. For example, Illumina sequencing platforms generally produce relatively short paired-end sequencing reads with high accuracy, while the more affordable Ion Torrent technology outputs single-end reads with often greater read lengths, but higher error rates (Quail et al., 2012; Fox et al., 2014; Salipante et al., 2014; Kwong et al., 2015; Escalona et al., 2016). Which sequencing platform different laboratories choose to acquire is not only dependent on financial resources, but also on individual needs and routine applications, with throughput, error rates / error types, read lengths and run time as the main concerning parameters. This leads to an increased diversification of the sequencing community (Moran-Gilad et al., 2015), creating a natural competition between producers, which benefits users through an ongoing improvement of technology and equipment. However, diversification also hampers standardization and despite ongoing calls for the establishment of agreed minimal sequencing quality parameters, this process has been much delayed (Endrullat et al., 2016).

Increasingly, microbial disease surveillance systems are based on WGS data. For example, Pathogenwatch (https://pathogen.watch) is a global platform for genomic surveillance, which analyses genomic data submitted by users and conducts cgMLST clustering to monitor the spread of important bacterial pathogens. Similarly, the GenomeTrakr network uses whole-genome sequence data and performs cg/wgMLST and SNP calling to track food-borne pathogens (https://www.fda.gov/food/whole-genome-sequencing-wgs-program/genometrakr-network) integrated into NCBI Pathogen Detection (https://www.ncbi.nlm.nih.gov/pathogens/). In Germany, a network of Federal State Laboratories and Federal Research Institutions supports the investigation of food-borne outbreaks through traditional typing and WGS methods. All genomic surveillance systems have in common that a high quality and accuracy of the sequencing data is crucial for a robust and reliable data analysis.

Proficiency testing (PT) is an important external quality assessment tool to ensure the accuracy and reproducibility of sequence data (Endrullat et al., 2016), whereas the aim of an interlaboratory study is to determine the variability of the results obtained by different collaborators. Several PT exercises with the focus on the sequencing of microbial pathogens have been published in recent years. In 2015, the GenomeTrakr network conducted a PT with 26 different US laboratories, which were instructed to sequence eight bacterial isolates according to a fixed protocol (Timme et al., 2018). In the same year, the Global Microbial Identifier (GM) initiative conducted an extensive survey with the aim to assess requirements and implementation strategies of PTs for bacterial whole genome sequencing (Moran-Gilad et al., 2015), followed by a series of global PT exercises (https://www.globalmicrobialidentifier.org/Workgroups/GMI-Proficiency-Test-Reports). In an interlaboratory exercise in 2016, five laboratories from three European countries (Denmark, Germany, the Netherlands) were asked to sequence 20 *Staphylococcus aureus* DNA samples according to a specific protocol and report cgMLST cluster types (Mellmann et al., 2017). In this study, we present the results of an interlaboratory study for short-read bacterial genome sequencing with ten participating laboratories from German-speaking countries initiated by the §64 German Food and Feed Code (LFGB) working group “NGS Bacterial Characterisation” chaired by the Federal Office of Consumer Protection and Food Safety (BVL). The working group serves to validate and standardize WGS methods for pathogen characterization in the context of outbreak investigations. The interlaboratory study was carried out by the German Federal Institute of Risk Assessment (BfR) in 2019, with the aim to answer the question whether different WGS technology platforms provide comparable sequence data, taking into account the routine sequencing procedures established in these laboratories.

## 2. Materials and Methods

### 2.1 Study design

In the frame of the §64 LFGB working group “NGS Bacterial Characterisation”, we conducted a interlaboratory study for next-generation sequencing. Twelve teams participated in the study. Participants included four Federal Research Institutes (3 German, 1 Austrian), four German State Laboratories, one German university and three German companies.

Participants were provided with DNA samples (40-55 μl, 60-187 ng/μl) of six bacterial isolates (Table 1) (two of each *Campylobacter jejuni, Listeria monocytogenes* and *Salmonella enterica*), with the species of the sample visibly marked on the tube containing the sample DNA.

**Table 1:** Strain characteristics of analysed DNA samples. (species, serovar, MLST, size and GC content) used in the interlaboratory study.

| Sample | Species | Serovar | MLST | Size (Mbp) | GC (%) |
| --- | --- | --- | --- | --- | --- |
| 19-RV1-P64-1 | *Campylobacter jejuni* |  | 4774 | 1.6 | 30.5 |
| 19-RV1-P64-2 | *Campylobacter jejuni* |  | 21 | 1.7 | 30.5 |
| 19-RV1-P64-3 | *Listeria monocytogenes* | IIc | 9 | 3.0 | 38.0 |
| 19-RV1-P64-4 | *Listeria monocytogenes* | IIb | 59 | 3.0 | 37.9 |
| 19-RV1-P64-5 | *Salmonella enterica* subsp. *enterica* | Infantis | 32 | 5.1 | 52.0 |
| 19-RV1-P64-6 | *Salmonella enterica* subsp. *enterica* | Paratyphi B var. Java | 28 | 4.8 | 52.2 |

Participants were instructed to sequence the samples according to their standard inhouse sequencing procedure. Where possible, participants were asked to sequence each isolate in two independent sequencing runs. No minimum quality criteria for the resulting sequencing data were requested. Together with the samples, participants received a questionnaire to document their applied sequencing method. Participants were given four weeks to conduct the sequencing and report the resulting raw sequencing data. Sequencing data was exchanged through a cloud-based platform and data quality was centrally analysed with open-source programs and in-house bioinformatic pipelines. Results of the sequencing data analysis were presented to the members of the §64 LFGB working group in November 2019. Following the meeting, ten participants agreed to a publication of the results of the interlaboratory study. Two participants declined a publication of their data due to a conflict of interest. Participants are anomalously identified with their laboratory code LC01 – LC10 assigned for this study.

### 2.2. Study isolates, cultivation and DNA isolation

Detailed information to the samples is summarized in Supplementary File 1 (Tables S1 to S3). The samples 19-RV1-P64-1 and 19-RV1-P64-2 were obtained from *Campylobacter jejuni* isolates (MLST type 4774 and 21 respectively). *Campylobacter jejuni* were precultured on Columbia blood agar, supplemented with 5 % sheep blood (Oxoid, Wesel, Germany) for 24 hours at 42 °C under micro-aerobic atmosphere (5% O_2_; 10% CO_2_). A single colony was inoculated on a fresh Columbia blood agar plate for an additional 24 hours. After incubation, bacterial cells were re-suspended in buffered peptone water (Merck, Darmstadt, Germany) to an OD600 of 2. Genomic DNA was extracted from this suspension with the PureLink^®^ Genomic DNA Mini Kit (Thermo Fisher Scientific, Dreieich, Germany) according to manual instructions.

The samples 19-RV1-P64-3 and 19-RV1-P64-4 were obtained from *Listeria monocytogenes* serovar IIc and serovar IIb respectively. *Listeria monocytogenes* were cultured on sheep blood agar plates and incubated at 37°C over night. Genomic DNA was directly extracted from bacterial colonies using the QIAamp DNA Mini Kit (Qiagen, Hilden, Germany) following the manual instructions for gram-positive bacteria.

The samples 19-RV1-P64-5 and 19-RV1-P64-6 were obtained from *Salmonella enterica* subsp. *enterica* serovar Infantis and serovar Paratyphi B var. Java respectively. *Salmonella enterica* were cultivated on LB agar (Merck). A single colony was inoculated in 4 ml liquid LB and cultivated under shaking conditions (180–220 rpm) at 37 °C for 16 hours. Genomic DNA was extracted from 1 ml liquid cultures using the PureLink® Genomic DNA Mini Kit (Thermo Fisher Scientific) according to manual instructions. DNA quality of all samples was verified with Nanodrop and Qubit and samples were stored at 4 °C before being express shipped in liquid form on ice.

### 2.3. PacBio reference sequences

As Pacific Biosciences (herein abbreviated as PacBio) sequencing was performed before the interlaboratory study started, DNA extractions used for PacBio sequencing differentiated from DNA extractions used for short read-sequencing. For *Campylobacter jejuni*, *Listeria monocytogenes* and *Salmonella enterica* the PureLink^®^ Genomic DNA Mini Kit (Invitrogen) was used for genomic DNA extraction.

PacBio sequences for samples 19-RV1-P64-1 to 19-RV1-P64-5 were obtained from GATC as described before (Borowiak et al., 2018).

Sample 19-RV1-P64-6 was sequenced in-house. Genomic DNA was sheared to approximately 10 kb using g-Tubes (Covaris, Brighton, U.K.) and library preparation was performed using the SMRTbell Template Prep Kit 1.0 and the Barcode Adapter Kit 8A (Pacific Bioscienses, Menlo Park, CA, USA). Sequencing was performed on a PacBio Sequel instrument using the Sequel® Binding Kit and Internal control Kit 3.0 and the Sequel® Sequencing Kit 3.0 (PacBio). Long read data was assembled using the HGAP4 assembler.

Information to the PacBio sequences is summarized in Supplementary File 1 (Tables S2-S3).

### 2.4. Whole-genome short read sequencing

All ten participants followed their own in-house standard protocol for sequencing. Important sequencing parameters such as the type of library preparation and sequencing kits, as well as the type of sequencing instrument were documented with a questionnaire (the questionnaire template in German language is provided as Supplementary File 2). The results of the questionnaire are summarized in Supplementary File 3. All participants determined the DNA concentration prior to sequencing library preparation. Of ten participants, seven chose a restriction digest for DNA fragmentation, while three laboratories fragmented DNA through mechanical breakage. Over half of participants pooled sequence libraries relative to genome sizes and almost all (with the exception of laboratory LC01) included a control in the sequencing run (i.e. PhiX).

All participants, with the exception of laboratories LC02 and LC08, sequenced samples in duplicates. Duplicates were defined as one sample sequenced in two independent sequencing runs on the same sequencing instrument, henceforth identified as sequencing run A and sequencing run B. Participants LC01, LC03, LC04, LC05, LC06, LC07, LC09, LC10 contributed 12 whole-genome sequencing data sets (combined forward and reverse reads) each, while participant LC08 contributed 6 whole-genome sequencing data sets. In contrast, laboratory LC02 sequenced the complete sample set on three different sequencing instruments in single runs, henceforth identified as LC02_a (Illumina iSeq), LC02_b (Illumina MiSeq), LC02_c (Illumina NextSeq). Therefore, participant LC02 contributed 18 whole-genome sequencing data sets.

Together, 120 whole-genome sequencing data sets were available for analysis. Taken the fact into consideration, that participant LC02 used three different sequencing instruments, a total of twelve individual sequencing instruments were included in the interlaboratory study: one Ion Torrent S5 instrument (Thermo Fisher Scientific), two iSeq, six MiSeq, two NextSeq and one NovaSeq instrument (all Illumina).

### 2.5. Assessment of raw sequencing data quality

The quality of the sequencing reads was assessed with fastp (Chen et al., 2018) with default parameters. Quality control parameters for each data set (forward and reverse reads for Illumina data) such as the number of total reads and the Q30 (before filtering) were parsed from the resulting fastp json reports. The coverage depth was calculated as the sum of the length of all reads divided by the length of the respective PacBio reference sequence.

### 2.6. Short-read genome assembling

Raw Ion Torrent reads were trimmed using fastp v0.19.5 (Chen et al., 2018) with parameters *--cut_by_quality3 --cut_by_quality5 --cut_window_size 4 -- cut_mean_quality 30*. Trimmed Ion Torrent reads were *de novo* assembled with SPAdes v3.13.1 (Nurk et al., 2013) with read correction.

Raw Illumina reads were trimmed and *de novo* assembled with our in-house developed Aquamis pipeline (https://gitlab.com/bfr_bioinformatics/AQUAMIS/) which implements fastp (Chen et al., 2018) for trimming and shovill (based on SPAdes) (https://github.com/tseemann/shovill) for assembly. Unlike SPAdes, shovill automatically down samples reads to a coverage depth of 100x prior to assembling.

### 2.7. Assessment of genome assembly quality and bacterial characterization

Quality of the genome assemblies was assessed with QUAST v5.0.2 (https://github.com/ablab/quast) without a reference. Quality parameters such as number of contigs, length of largest contig and N50 were parsed from the QUAST report text files for each assembly.

Based on the genome assemblies (including the PacBio reference sequences), bacterial characterization was conducted with our in-house developed Bakcharak pipeline (https://gitlab.com/bfr_bioinformatics/bakcharak) which implements among other tools, ABRicate for antimicrobial resistance and virulence factor screening (https://github.com/tsmeeann/abricate), and the PlasmidFinder database for plasmid detection (Carattoli et al., 2014), mlst (https://github.com/tseemann/mlst), SISTR (Yoshida et al., 2016) for *in silico* Salmonella serotyping and Prokka (Seemann, 2014) for gene annotation.

### 2.8. CgMLST allele calling

CgMLST allele calling was conducted with our in-house developed chewieSnake pipeline (https://gitlab.com/bfr_bioinformatics/chewieSnake) which implements chewBBACA (Silva et al., 2018). Only complete coding DNA sequences, with start and stop codon, according to the NCBI genetic code table 11, are identified as alleles by chewBBACA (with Prodigal 2.6.0 (Hyatt et al., 2010)). CgMLST allele distance matrices are computed with grapetree (ignoring missing data in pairwise comparison).

CgMLST schemes for *Listeria monocytogenes* (Ruppitsch et al., 2015) were derived from the cgMLST.org nomenclature server (https://www.cgmlst.org/). CgMLST schemes for *Campylobacter jejuni* and *Salmonella enterica* were derived from the chewBBACA nomenclature server (http://chewbbaca.online/).

### 2.9. SNP calling

SNP (single-nucleotide polymorphism) calling was conducted for each sample. Sequencing reads were trimmed prior to SNP calling. Assembled uncirculated PacBio sequences of the samples were used as reference sequences for SNP calling. SNP calling was conducted with our in-house developed snippySnake pipeline (https://gitlab.com/bfr_bioinformatics/snippy-snake) which implements snippy v4.1.0 (https://github.com/tseemann/snippy).

## 3. Results

### 3.1. Comparison of quality of sequencing reads

One important parameter to assess the quality of sequencing reads is the phred quality score. Commonly a Q score of 30 is used, which indicates a base call accuracy of ?99.9 %. We compared the percentages of bases that have a quality score equal or larger to 30. The results visualized in Figure 1 (see Supplementary File 4 for exact numbers), show that on average ~ 90 % of Illumina bases have a Q score ≥Q30, while only ~40 % of Ion Torrent bases achieve a Q score ≥Q30. Therefore, the base call accuracy of Illumina data is greater than that of Ion Torrent data. There is little variation within the Illumina instrument series (mean values: iSeq: 91.7 %; MiSeq: 90.8 %; NextSeq: 90.4 %; NovaSeq: 92.4 %), indicating that no particular instrument of the series out or under performs the others. In contrast, sequencing data with higher or lower quality scores was consistently associated with individual laboratories. Among the participants employing Illumina instruments, LC08 overall produced the lowest quality data (LC08 mean: 82.1 %), while LC02_b produced the highest quality data (LC02_b mean: 97.9 %), both with a MiSeq instrument. Interestingly, the same laboratory LC02 remained behind the average for Illumina data when employing a NextSeq instrument (LC02_c mean: 87.1 %). Of course, sequence quality might also depend on loading concentration and number of cycles used for sequencing. Quality scores remained largely consistent between runs. Equally, the type of bacterial species had little influence on sequencing data quality.

**Figure 1:**
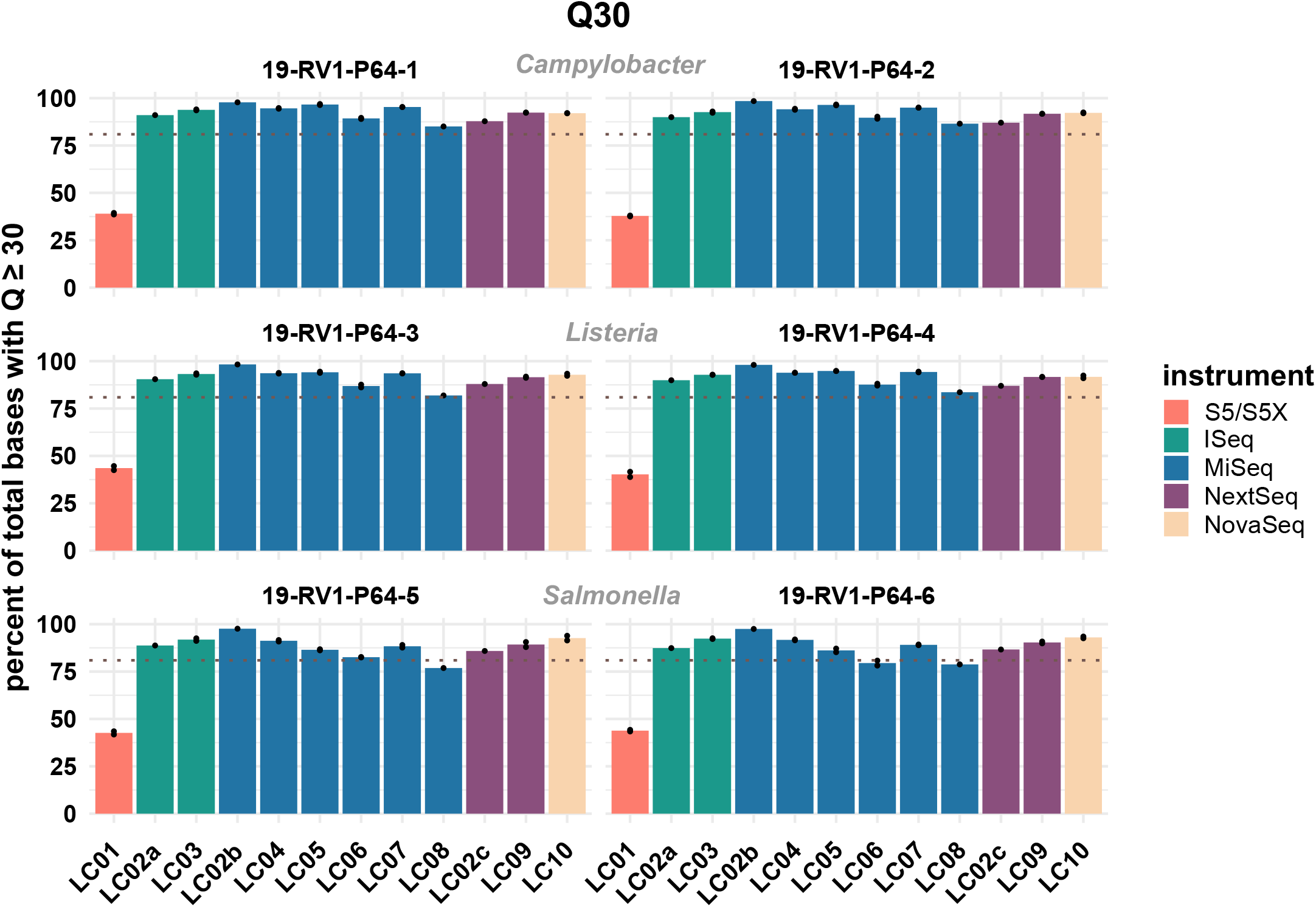
The bar plot shows the mean percentage of total bases with a phred score above or equal to Q30 grouped by laboratories and samples. Line-connected points indicate the variance between sequencing runs (run A / run B), with the exception of laboratories L02 and LC08 (single sequencing run). Fill colours identify the sequencing instrument. The species of the samples is indicated. The dotted line marks a Q30 of 80%.

We further assessed the total number of reads and bases of each data set. Since ideally there is little variation in the length of the reads (for Illumina), the number of reads is closely correlated with the total number of produced bases, as can be seen from Supplementary File 4. To achieve a reasonable coverage over the whole genome a minimum number of reads / total bases is required (this can be easily calculated when a suitable reference genome is available).

As visualized in Figure 2, the total number of produced bases varied across laboratories, instruments and samples, as well as between sequencing runs. For example, for sample 19-RV1-P64-1, laboratory LC10 produced the greatest number of sequencing bases: ~1.8 billion base (~12.2 million reads), while laboratory LC02_b produced the lowest number of sequencing bases: ~0.8 billion bases (~0.4 million reads). The number of reads / total number of bases has a direct influence on the coverage depth (in this study calculated by the total number of bases divided by the length of the PacBio reference). Sufficient coverage depth is an important requirement for successful downstream analysis, such as variant detection and assembly. However, up to now there is no widespread consensus for the recommended minimum coverage depth for bacterial whole genome sequencing. In the accompanying questionnaire, participants stated that they intended to achieve a coverage depth ranging from >20x to <300x, with most participants opting for a coverage depth of 60x to 70x. Actual coverage depths ranged from 26x (LC03, 19-RV1-P64-5, run A) to 1201x (LC10, 19-RV1-P64-1, run B), with most data sets featuring coverage depths from 75x to 196x (Q_0,25_ and Q_0,75_). With the exception of a small number of data sets (LC03: 19-RV1-P64-2, 19-RV1-P64-5, 19-RV1-P64-6; LC05: 19-RV1-P64-6, all run A), all other data sets were well above a coverage depth of 30x. Similarly, to the total number of produced reads, actual coverage depths varied between laboratories, instruments and samples, as well as between sequencing runs. In concordance with the high number of total reads / bases, laboratory LC10 produced data sets with very high coverage depths with an average of 736x. When coverage depths were normalized, by assigning a coverage depth of 1 to sample 19-RV1-P64-1 of each group, we found that coverage depths varied in a predictable manner in relation to the genome size of the sample as shown in Figure 4. Some participants chose to pool sequencing libraries relative to genome sizes of the samples, which in most cases ensured a more consistent sequencing depth across the samples (LC02_a, LC03, LC04, LC06). In comparison, participants that pooled sequencing libraries of all samples equally (LC01, LC05, LC07, LC08, LC10) obtained lower coverage depths for bacterial isolates with larger genome sizes (i.e. ~4.9 Mbp for *Salmonella enterica*), and high coverage depths for bacterial isolates with smaller genome sizes (i.e. ~1.7 Mbp *Campylobacter jejuni*). However, in most cases pooling the DNA libraries relative to genome size only reduced the impact of the genome size effect, without eliminating it. Only laboratory LC06 achieved a high consistency across all samples.

**Figure 2:**
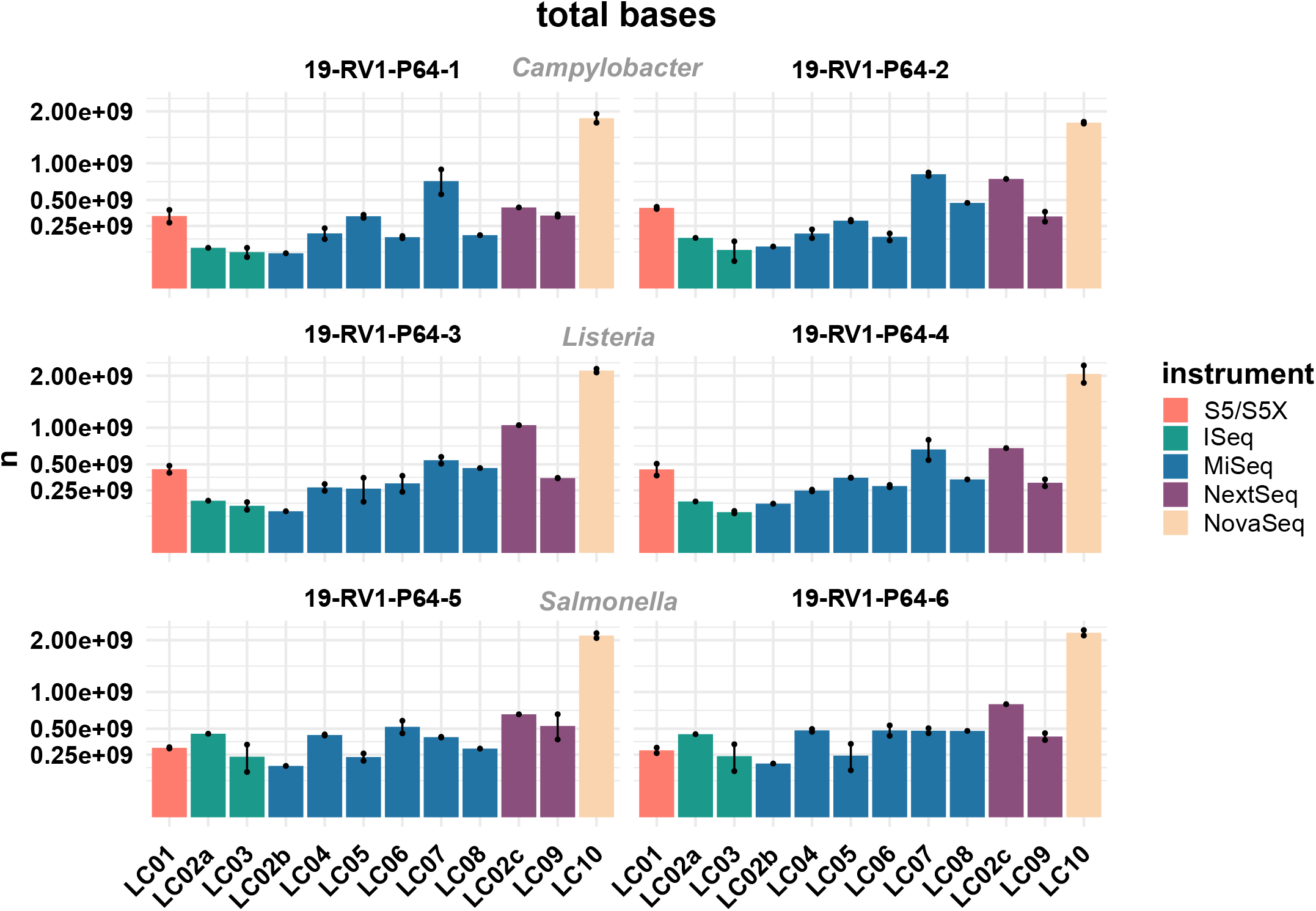
The bar plot shows the mean total number of bases encompassed by the raw reads grouped by laboratories and samples. Line-connected points indicate the variance between sequencing runs (run A / run B), with the exception of laboratories L02 and LC08 (single sequencing run). Fill colours identify the sequencing instrument. The species of the samples is indicated. The y-axis is squared.

**Figure 3:**
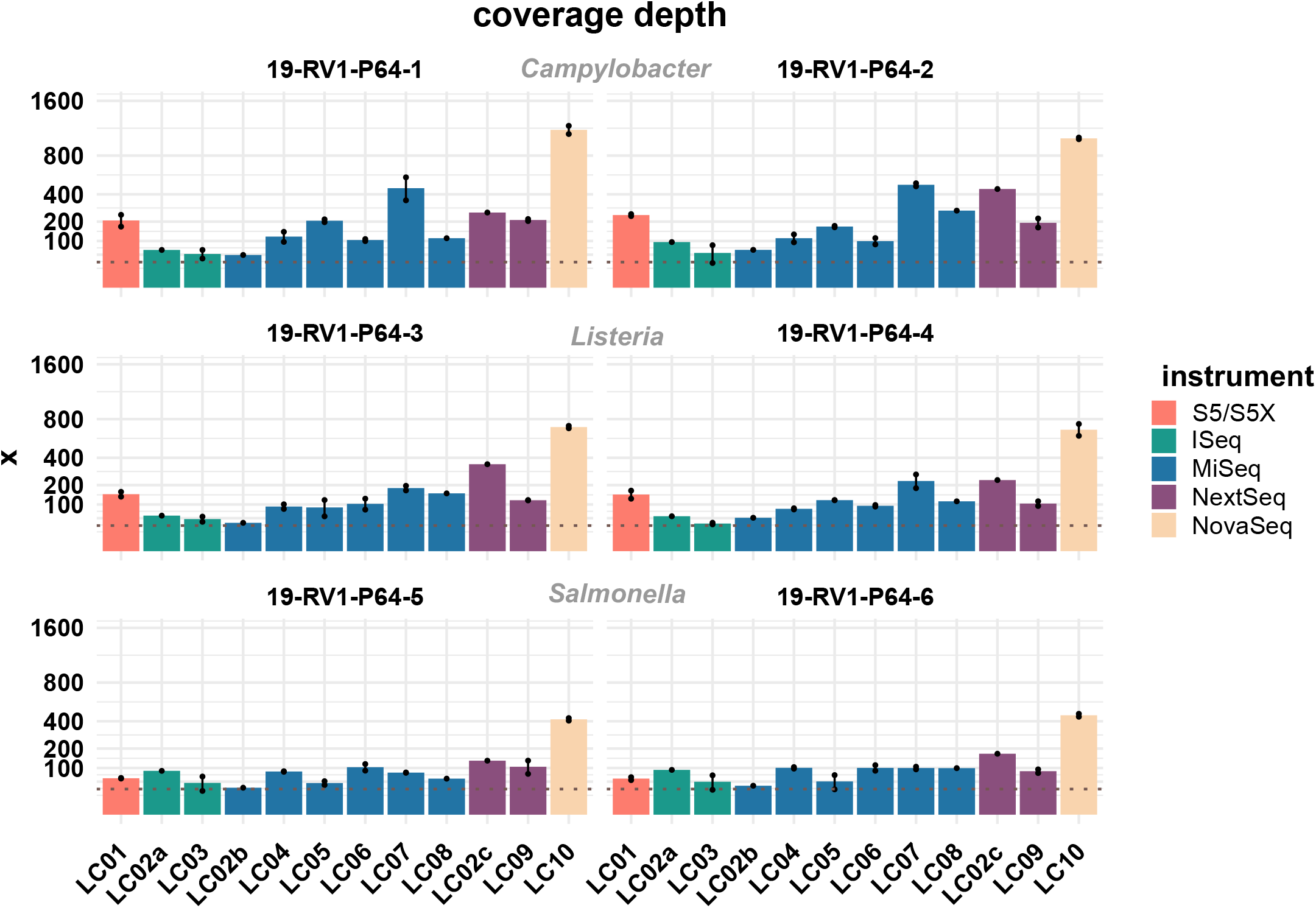
The bar plot shows the mean coverage depth grouped by laboratories and samples. Line-connected points indicate the variance between sequencing runs (run A / run B), with the exception of laboratories L02 and LC08 (single sequencing run). The coverage depth was defined as the sum of the length of all raw reads divided by the length of the respective PacBio reference sequence. Fill colours identify the sequencing instrument. The species of the samples is indicated. The y-axis is squared. The dotted line marks a coverage depth of 30x.

**Figure 4:**
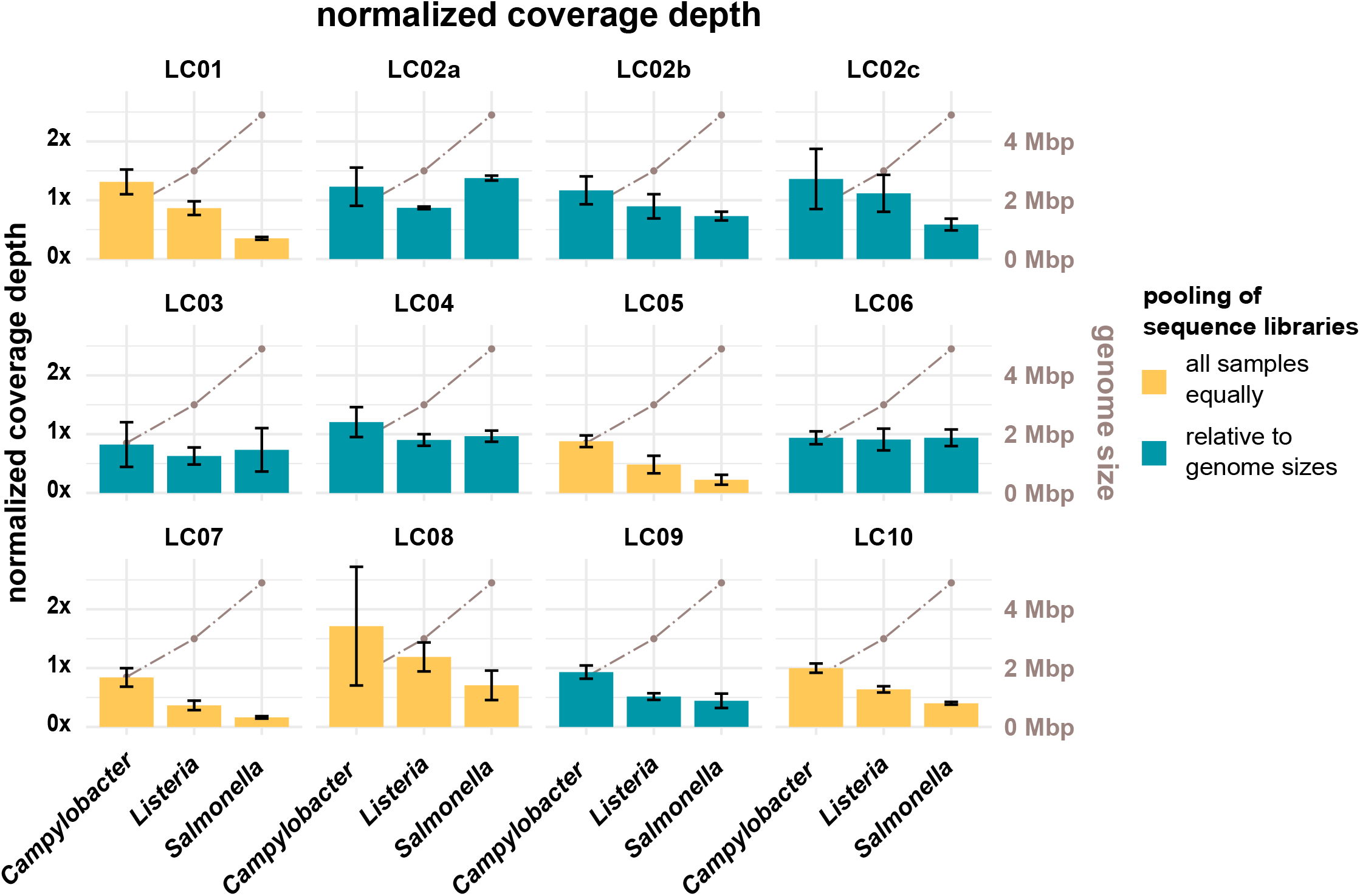
The bar plot (left y-axis) shows the mean normalized coverage depth grouped by laboratories and species of the samples with error bar. The coverage depth was normalized for each laboratory to the coverage depth for sample 19-RV1-P64-1, sequencing run A, which was assigned a value of 1. The coverage depth was defined as the sum of the length of all raw reads divided by the length of the respective PacBio reference sequence. Fill colours identify, whether DNA libraries were pooled relative to genome sizes prior to sequencing or whether DNA libraries were pooled equally. The brown line graph in the background (right y-axis) indicates the average genome size of the species.

### 3.1. Comparison of quality of genome assemblies and bacterial characterization

The genome assemblies constructed from the short read data were assessed and all determined quality parameters are listed in Supplementary File 4. We found little variation in the length of the genome assemblies within the short read assemblies (sd values for the samples ranged from ~3 Kbp to ~11 Kbp). However, all short read assemblies were ~36 to ~66 Kbp shorter than their respective PacBio references, likely due to overlapping end regions in the PacBio sequences, which were not circularized prior to analysis.

Similarly, there was little variation for the calculated GC values (sd values for the samples ranged from 0.01 to 0.03 %). Besides the length, the quality of genome assemblies is determined by the total number of contigs, and the size of the largest contig, with assemblies featuring fewer, larger contigs generally being more useful for downstream analyses. Both parameters are combined in the N50 value, which is defined as the length of the shortest contig in the set of largest contigs that together constitute at least half of the total assembly size. The N50 values for all assemblies are visualized in Figure 5. We found N50 values to be overall very similar for individual samples, regardless of which laboratory or instrument provided the sequencing data, with a few notable exceptions (i.e. LC06, LC08). In general, highest N50 values were obtained for *Listeria monocytogenes* samples (19-RV1-P64-3: ~600 Kbp; 19-RV1-P64-4: ~480 Kbp), followed by *Salmonella enterica* samples (19-RV1-P64-5: ~200 Kbp; 19-RV1-P64-6: ~340 Kbp), and *Campylobacter jejuni* samples (19-RV1-P64-1: ~220 Kbp; 19-RV1-P64-2: ~180 Kbp).

**Figure 5:**
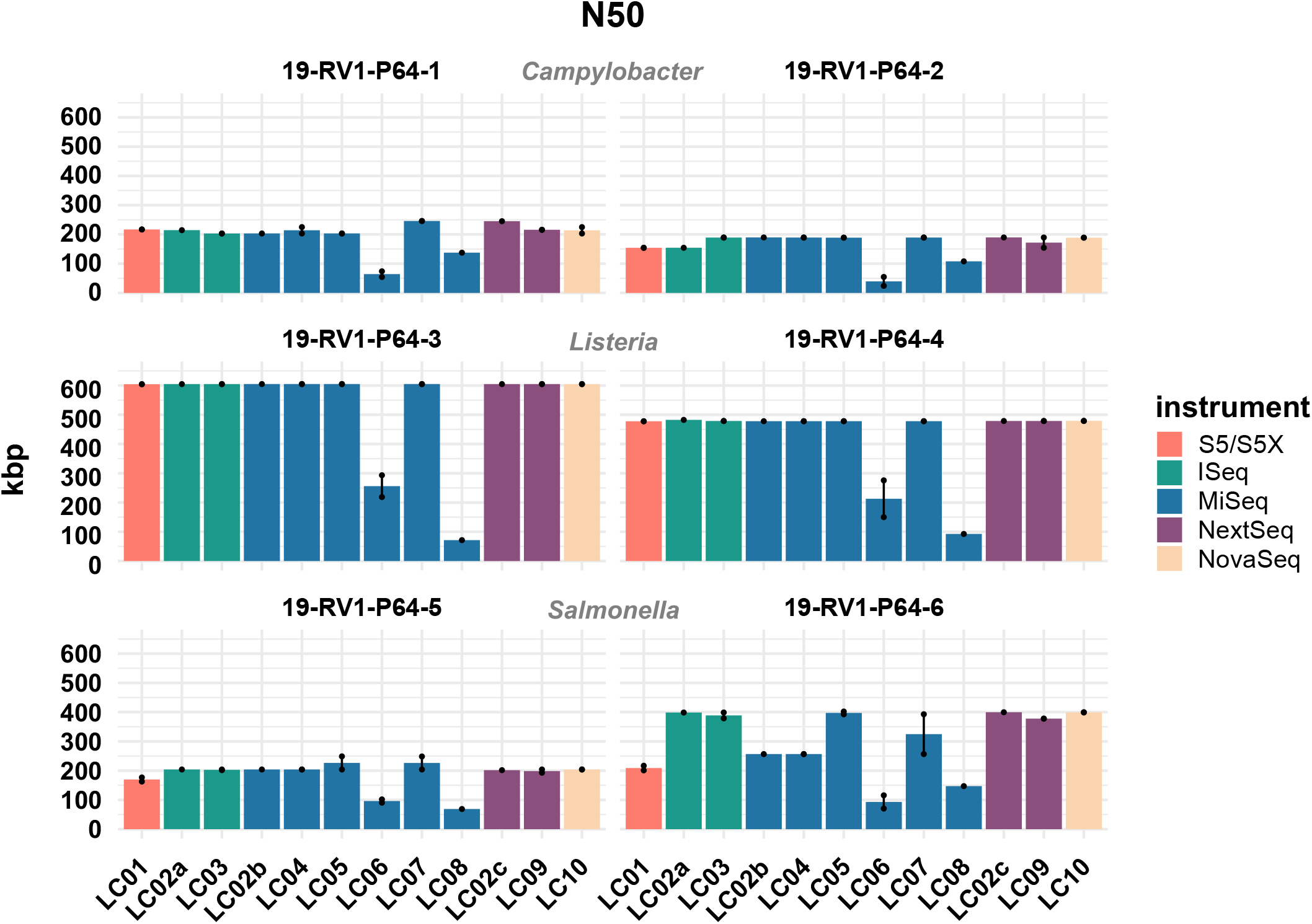
The bar plot shows the mean N50 determined for the short-read genome assemblies grouped by laboratories and samples. Line-connected points indicate the variance between sequencing runs (run A / run B), with the exception of laboratories L02 and LC08 (single sequencing run). Fill colours identify the sequencing instrument. The species of the samples is indicated.

Assemblies of laboratories LC06 and LC08 consistently had much lower N50 values (also shown by a higher total number of contigs and shorter contigs lengths), compared to the rest of the group. For example, while the majority of assemblies achieved an N50 of ~ 605 Kbp (± 550 bp) for sample 19-RV1-P64-3, the N50 for assemblies of LC06 ranged around ~ 256 Kbp, while the N50 for assemblies of laboratory LC08 was even lower (~71 Kbp). Interestingly, no linear correlation was apparent between the N50 value and the coverage depth as shown in Figure 6.

**Figure 6:**
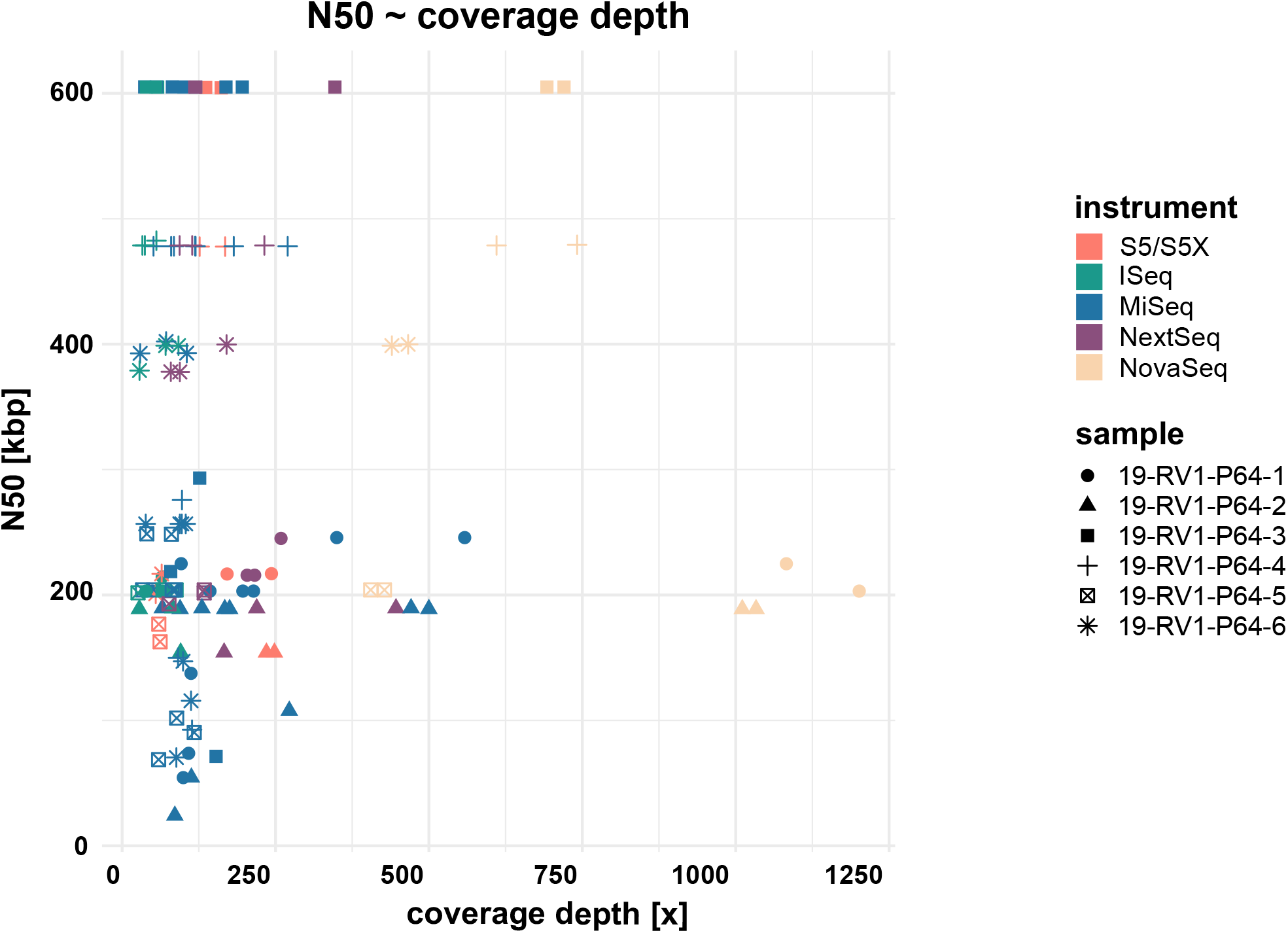
The dot plot shows the correlation between N50 and coverage depth for the short-read genome assemblies / sequence data sets. Fill colours indicate the sequencing instrument.

Coding frames in the genome assemblies were annotated to determine the MLST type, as well as resistance and virulence genes. In total, there was little variation for the total number of detected CDS (defined as a sequence containing a start and stop codon). The total number of CDSs varied by sample (19-RV1-P64-1: n=~1597; 19-RV1-P64-2: n=~1713; 19-RV1-P64-3: n=~2892; 19-RV1-P64-4: n=~2913; 19-RV1-P64-5: n=~4667; 19-RV1-P64-6: n=~4393) with a standard deviation of 8 to 15 coding frames.

The Multilocus Sequence Type (MLST) was determined correctly for all data sets. The same plasmid markers could be detected from all short read genome assemblies. Two more plasmid markers (*Col8282_1* and *ColRNAI_1*) could be detected in the short read assemblies compared to the PacBio reference for 19-RV1-P64-6, likely due to the fact that small plasmids are often excluded from PacBio sequences (read lengths too short). In three cases, resistance genes detected in the PacBio references were not present in the short read assemblies: *bla*_OXA-184_ in 19-RV1-P64-1, of laboratory LC06 (run A) and *aadA1* in 19-RV1-P64-6, of laboratory LC09 (both runs).

Although overall the same virulence genes could be detected from all short-read assemblies, there was some variation with assemblies from laboratories LC01, LC06 and LC08 often missing virulence genes (Supplementary File 4). For example, virulence factors *flaA* and *flaB* could not be detected in assemblies from laboratory LC01 for sample 19-RV1-P64-1. Interestingly, the same two genes were present in both assemblies of laboratory LC01 for sample 19-RV1-P64-2, but absent in all other assemblies for this sample. In another example the genes *sopD2* and *sseK1* could not be detected from the assembly for sample 19-RV1-P64-5 from laboratory LC08. The absence of virulence and resistance genes is likely caused by contig borders.

### 3.3. CgMLST calling

CgMLST was conducted to compare the effect of differences in the genome assemblies on clustering. All cgMLST distance allele matrices are presented in Supplementary File 5. The cgMLST distance matrix for sample 19-RV1-P64-1 is visualized in Figure 7. CgMLST distance matrices for the six samples were overall very similar. In general, most assemblies had zero allele differences. However, assemblies constructed from Ion Torrent short read data (LC01) generally had a much higher number of allele differences, than those constructed from Illumina short reads. For easy comparison, we calculated the ‘median cgMLST distance’ for each assembly, by computing the medium of all allele differences to a specific assembly (compare Figure 7).

**Figure 7:**
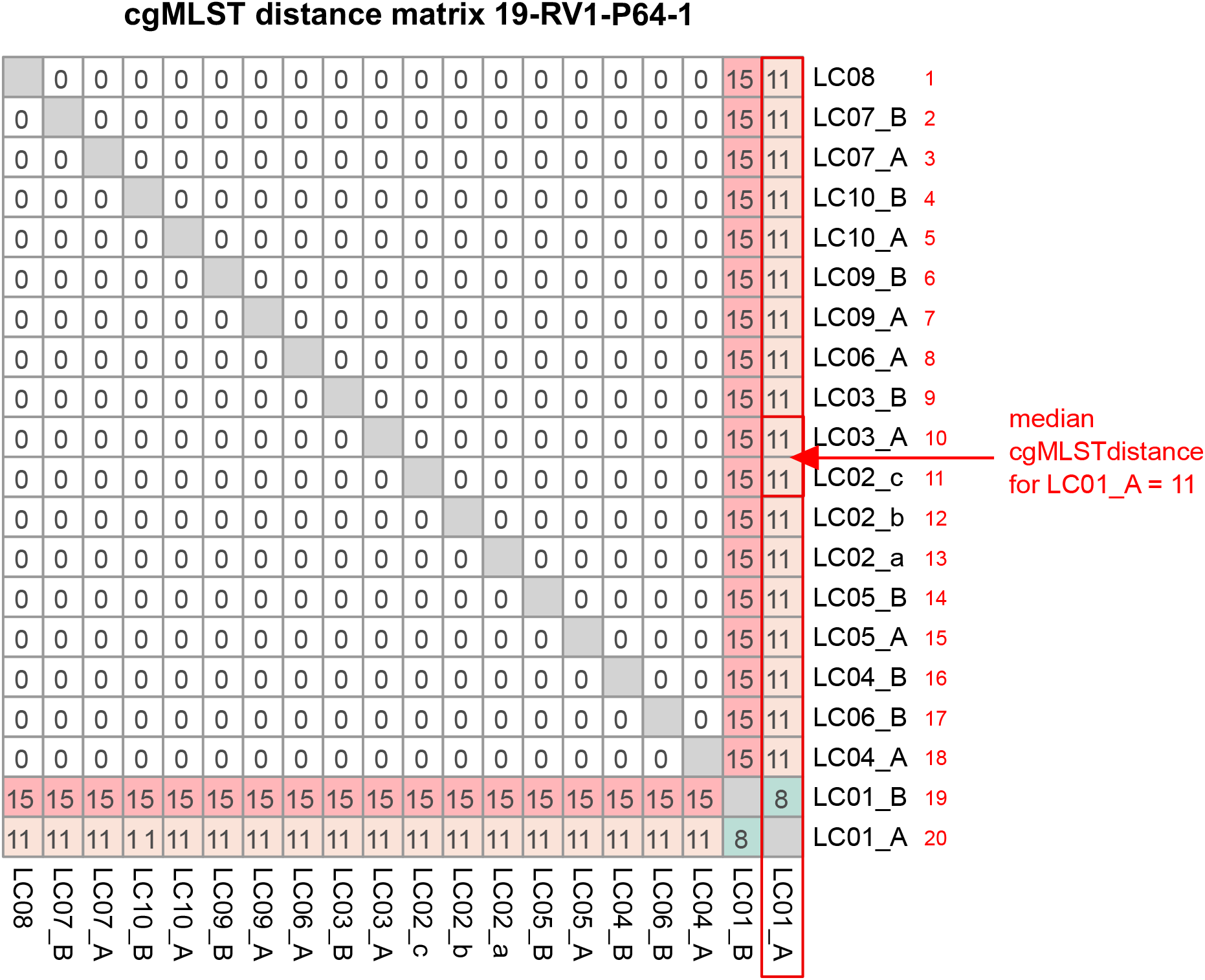
The figure shows the cgMLST distance matrix for sample 19-RV1-P64-1. Laboratories (LC01-LC10) and respective sequencing runs (run A / run B) are identified. The red box, arrow and text demonstrate how the median cgMLST distance was determined.

Figure 8 shows the median cgMLST distance for all assemblies. As mentioned the highest number of allele differences were calculated for the assemblies of laboratory LC01 (using an Ion Torrent instrument). However, allele differences for the Ion Torrent assemblies varied dependent on the species of the sample. The smallest number of cgMLST allele differences were obtained for *Listeria monocytogenes* samples (LC01: ~ 7.1), followed by *Campylobacter jejuni* samples (LC01: ~ 11.1) and *Salmonella enterica* samples (LC01: ~ 26.1). Illumina assembly generally had much lower allele differences. Median cgMLST allele differences for the assemblies of the laboratories LC02a, LC02b, LC02c, LC03, LC04, and LC010 were zero for all samples. Median allele differences for assemblies of the laboratories LC05, LC06, LC07, LC08, and LC09 were between zero and three, often slightly higher for laboratories LC05 and LC08. Interestingly, the assembly of sample 19-RV1-P64-6 produced in run A by LC05 featured a median number of 10 alleles, while the assembly produced in the independent run B by LC05 had a median number of zero allele differences.

**Figure 8:**
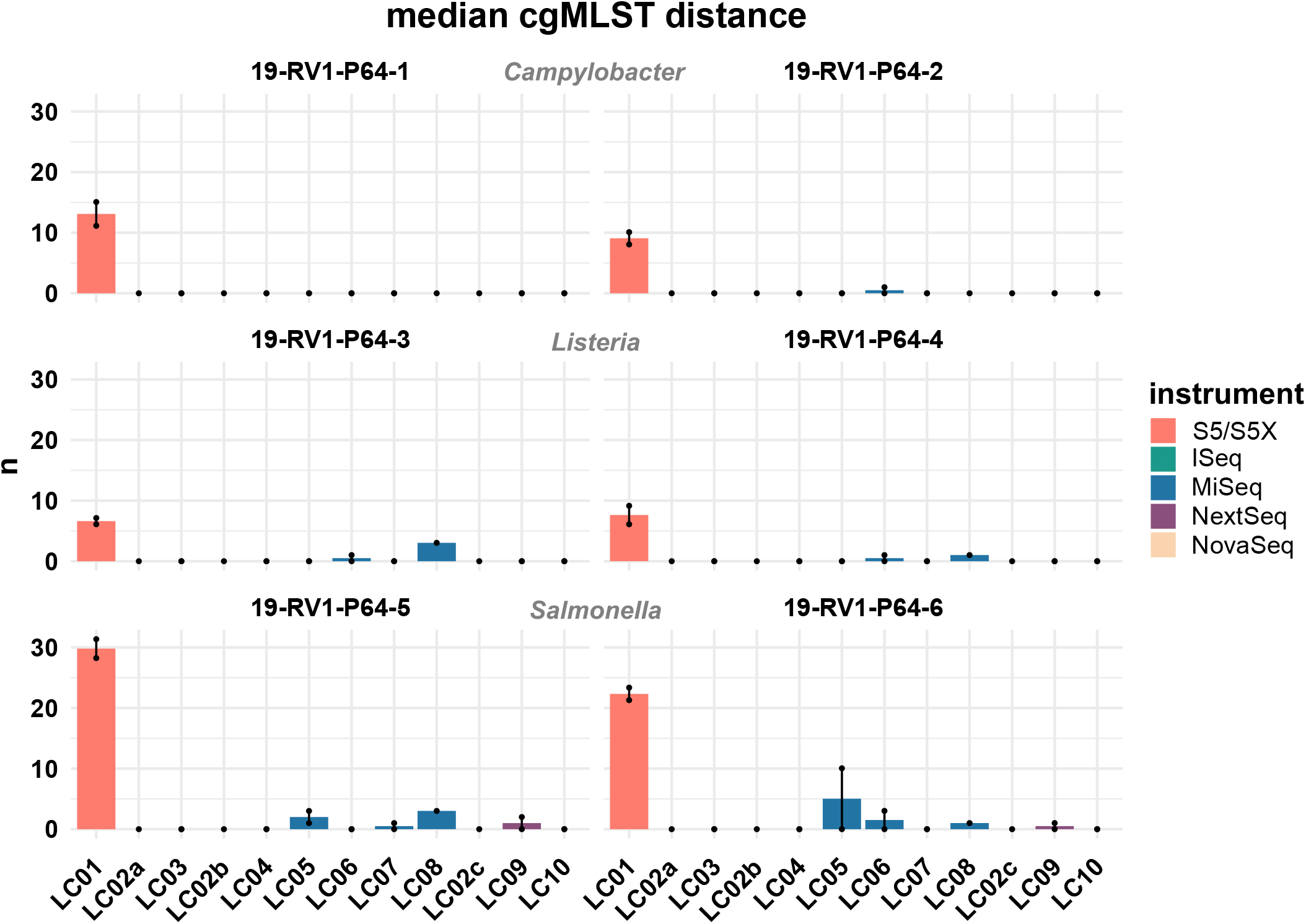
The bar plot shows the mean median cgMLST distance grouped by laboratories and samples. Line-connected points indicate the variance between sequencing runs (run A / run B), with the exception of laboratories L02 and LC08 (single sequencing run). Fill colours identify the sequencing instrument. The species of the samples is indicated.

We further compared the effect of the assembly algorithm on the cgMLST calling by assembling trimmed Illumina reads with SPAdes (as opposed to shovill) prior to cgMLST calling. However, no significant difference was found in the number of allele differences (data not shown).

### 3.4. SNP calling

SNP calling was conducted to detect sequencing errors. The assembled PacBio sequences were used as reference sequences. All SNP distance allele matrices are presented in Supplementary File 6. No SNPs were detected within the data sets. Equally, all data sets featured zero SNPs to the reference sequence, with the exception of the PacBio reference for sample 19-RV1-P64-5, to which all data sets had 2 SNPs.

## Discussion

We conducted an interlaboratory study for the investigation of the reproducibility and consistency of bacterial whole-genome sequencing. Ten participants were instructed to sequence six DNA samples in duplicate according to their in-house standard procedure protocol. We were interested to see, how the quality of sequencing data varied across different sequencing instruments, library preparation kits, sequencing kits and individual expertise of the participating laboratories. Overall, we were able to compare 12 Illumina sequencing instruments and one Ion Torrent instrument.

It is well known that different sequencing technologies vary in their average error rates, with Ion Torrent data generally having higher error rates compared to Illumina (Quail et al., 2012; Fox et al., 2014; Salipante et al., 2014; Kwong et al., 2015; Escalona et al., 2016). Indeed, we assessed that Ion Torrent bases achieved much lower quality scores than Illumina bases (Ion Torrent Q30: ~ 35-50 %, Illumina Q30: ~ 80 – 95 %). Interestingly, we found the four different Illumina sequencing instruments types involved in our study (iSeq, MiSeq, NextSeq, NovaSeq) to be very similar in terms of base quality, suggesting that the underlying sequencing technology is very similar, despite the different color chemistry used.

There was a great variety in the number of total bases that participants obtained for their data sets, resulting in great fluctuations for the coverage depth (ranging from 26x to 1200x). Although no widely accepted minimal coverage depth for bacterial wholegenome sequencing is established yet, most studies recommended coverage depths ranging from ≥30x to ≥50x (Chun et al., 2018). Positively, most data sets submitted by the participants in our study had coverage depths well above 30x, demonstrating that insufficient coverage depth is not usually a concern. However, coverage depths frequently fell short of the intended coverage depths stated by participants in the accompanying questionnaire, indicating that this parameter is not always well controlled for. For example, while laboratory LC02_b aimed for a coverage depth of ≥60x, the majority of data sets submitted by this laboratory had a much lower coverage depth (30-50x). Similarly, laboratory LC01, LC02a, LC05 and LC08 frequently obtained lower than intended coverage depths.

Resulting from experience and producer instructions, users generally know the number of reads / total bases that their sequencing instrument is capable of producing in one sequencing run. By pooling DNA libraries relative to genome sizes (provided the species of the isolates is known), users can influence the number of reads / bases and therefore the coverage depth for each isolate. As was shown in this study, participants that pooled DNA libraries prior to sequencing relative to genome sizes achieved more consistent coverage depths across the three species (e.g. LC06), while participants that pooled all DNA libraries equally, obtained sequencing data with predictable fluctuation in coverage depth (i.e. LC10), depending on the genome size of the organism.

Both, too low (problematic for variance calling / fragmented assembly) and too high (increased ‘noisiness’ of the data since the number of sequencing errors increases with the read number / the assembly graph is too complex and cannot be resolved) coverage depths can have negative effects on downstream analysis. For this reason, updated assembly algorithm, such as shovill, ‘down sample’ data to a moderate coverage prior to assembly (e.g. shovill down samples to 100x). Indeed, we did not find a linear correlation between coverage depth and N50 (i.e. the very high coverage depths observed for some data sets had neither positive nor negative effects on assembly quality). Nevertheless, we recommend that sequencing laboratories pool DNA libraries by genome sizes prior to sequencing in order to produce sequencing data with consistent coverage depth for optimal downstream analysis. This has the additional benefit that smart pooling strategies decrease the sequencing costs, as a greater number of samples can be sequenced in one run.

We employed SNP calling for the detection of potential sequencing errors in the trimmed sequence reads, as well as for assessing the utility of a SNP calling approach for an integrated outbreak analysis with data from different sequencing platforms. Given that participants were provided with purified DNA samples, thereby eliminating the potential for the development of mutations during cultivation, any SNP potentially flags a sequencing error. Positively, we detected zero SNPs within the data sets. The fact that all data sets of sample 19-RV1-P64-5 differed in two SNPs from the respective PacBio reference, either points to a sequence error within the PacBio reference, or might indicate that the strain underwent mutations between the independent cultivations for short read and long read sequencing DNA isolation.

We further constructed *de novo* assemblies from the short read sequence data to assess the influence of variations in sequence data quality on assembly-based downstream analysis. To eliminate assembler specific effects we strove to construct all assemblies in an equal manner. Naturally, single-end Ion Torrent data requires different assembly algorithm, than those employed for paired-end Illumina data, which hampers a direct comparison.

Nevertheless, we found that all assemblies were overall very similar, with respect to assembly length, N50, GC and the number of CDSs, with a few notable exceptions. In particular, assemblies constructed from short read data of laboratories LC06 and LC08 (both using a MiSeq Illumina instrument) had much lower N50 values and a greater number of contigs, probably due their use of the Nextera XT DNA Library Preparation Kit, which was recently shown to have a strong GC bias (Grützke et al., 2019; Sato et al., 2019; Uelze et al., 2019) (also compare Supplementary File 7). This is a concern since a high number of contigs in a genome assembly may cause a fragmentation of genes at the contigs borders, thereby affecting gene annotation and multilocus sequence typing. Furthermore we found that Ion Torrent assemblies differed from Illumina assemblies in length (slightly shorter), N50 (slightly lower), GC (slightly lower) and number of CDSs (slightly increased).

Complementary to SNP calling, we employed a cgMLST approach to compare genome assemblies in a simulated outbreak analysis. Noteworthy, cgMLST revealed a major distinction between Illumina and Ion Torrent data with assemblies constructed from Ion Torrent reads generally computing a much greater number of allele differences (Illumina: ~ 0-3 allele differences, Ion Torrent: ~10-30 allele differences). We suspect that this increased number of allele differences is caused by frame shifts in the Ion

Torrent assemblies. While the typical error type associated with Illumina reads are randomly distributed incorrect bases (substitution error) which do not cause frame shifts, Ion Torrent reads are prone to systematic insertions and deletions errors which lead to frame shifts in coding sequences (Buermans and den Dunnen, 2014; Escalona et al., 2016). Given that the cgMLST method employed in this study identifies coding frames based on their start and stop codons (as opposed to methods which implement a similarity based BLASTn search against a set of reference loci for allele identification), frame shifts will have a major effect on allele detection, thereby likely causing the observed increased number of allele differences. This is further supported by the low reproducibility of the Ion Torrent assemblies with up to 24 allele differences between two independent sequencing runs for the same sample.

From our results, SNP calling seems to be the method of choice for a combined outbreak analysis which integrates Illumina and Ion Torrent data sets in concordance with earlier studies (Kaas et al., 2014), due to the fact that Ion Torrent typical indels, as well as heterozygous or low quality sites are excluded from SNP calling. Through SNP calling it was possible to correctly identify the clonality between data sets for the same sample (i.e. there were zero SNPs between the Illumina and the Ion Torrent data sets for all samples). CgMLST calling, on the other hand would have produced much confusion in a real outbreak study, by suggesting that DNA samples sequenced with Ion Torrent were obtained from isolates relatively unrelated from those sequenced with Illumina.

These seemingly contradictory results can be explained by the stringent variant filtering prior to SNP calling, which eliminates the effect of Ion Torrent typical insertion and deletion errors. However, masking of indels and other low quality sites might also decrease the number of SNPs detected in total, thus leading to a lower resolution. SNP calling further has the advantage that no assembling step is required, for which currently no optimized assembly algorithm is available for Ion Torrent, thereby avoiding the introduction of assembly biases. Although we additionally assembled Illumina reads with SPAdes to increase the comparability to Ion Torrent assemblies (currently shovill is unable to assemble Ion Torrent reads), SPAdes remains inherently tailored for Illumina reads and cgMLST calling was not improved with all SPAdes assemblies. Given that many surveillance platforms perform cgMLST or wgMLST for (pre-)clustering (Uelze et al., 2020) the observed differences between Illumina and Ion Torrent assemblies might potentially lead to erroneous clustering results and disrupt outbreak studies.

## Conclusion

We found that seven of nine participants with Illumina sequencing instruments were able to obtain reproducible sequence data with consistent high quality. Two participants with Illumina instruments submitted data with lower quality, probably due to the use of a library preparation kit, which shows difficulty in sequencing low GC genome regions. The only Ion Torrent instrument included in our study was inferior in terms of sequence data quality and assembly accuracy. We found a SNP calling approach to be more suitable for an integrated data and outbreak analysis of Ion Torrent and Illumina data than a cgMLST calling approach.

In the future, sequencing laboratories will continue to adapt and modify their laboratory protocols in order to optimize sequencing data quality, throughput and user-friendliness, while striving for the most cost and time-effective procedure. We welcome these efforts by innovative and thoughtful staff, which should not be unnecessarily hampered by overly rigid procedural protocols. Instead, a set of widely accepted, scientifically based and sensible minimal sequencing quality parameters, together with good standard practice protocols are urgently needed to ensure a consistent high quality of sequencing data for comparative data analysis.

Continuous interlaboratory testing, such as the one employed in this study and external PTs, will play an important role in ensuring that laboratories of the diverse public health setting adhere to these standards, while providing important feedback to participants on their competency level. Open or anonymous sharing of sequencing parameters allows an assessment of the utility of different sequencing approaches and helps to identify potential user issues. In the best case, interlaboratory studies promote knowledge and expertise sharing, enabling laboratories to adopt the sequencing procedures best suited for their unique setting, while simultaneously contributing to a standardization of the technology, which will greatly improve the efficacy of sequencing data for surveillance, outbreak analyses and comparative studies.

## Abbreviations

BLAST: basic local alignment search tool
cgMLST: core genome multilocus sequence typing
DNA: deoxyribonucleic acid
MLST: multilocus sequence typing
NGS: next-generation sequencing
SNP: single-nucleotide polymorphism
ST: sequence type
PT: Proficiency testing
wgMLST: whole-genome MLST
WGS: whole genome sequencing

## Data Availability

Sequencing data for all data sets analysed in this study has been deposited in the European Nucleotide Archive (ENA) under the study accession number PRJEB37768.

## Acknowledgements

This work was supported by the German Federal Institute for Risk Assessment (BfR).

The BfR has received financial support from the Federal Government for Laura Uelze on the basis of a resolution of the German Bundestag by the Federal Government and funded by the Ministry of Health within the framework of the project “Integrated genomebased surveillance of *Salmonella* (GenoSalmSurv)”, decision ZMVI1-2518FSB709 of 26/11/2018.

LU, MB and BM designed the study. LU and MB coordinated the interlaboratory study. LU and CD conducted the bioinformatic analysis and evaluation of the sequencing data quality. CD and ST developed the in-house bioinformatic pipelines used for analysis of the sequencing data. BM supervised the project. LU wrote the manuscript and created the figures. All authors read and approved the manuscript.

We are grateful for the continuous collaboration with the National Reference Laboratory for *Salmonella*, as well as the National Reference Laboratory for *Listeria monocytogenes* and the National Reference Laboratory for *Campylobacter* who kindly provided us with the bacterial isolates and DNA samples.

We thank Beatrice Baumann and Katharina Thomas (BfR), Sara Walter (LAVES) and Adrian Prager (MRI) for their excellent laboratory assistance.

## Supplementary Files

Supplementary File 1:

**Table S1:** General information about the strains used for the interlaboratory study.

**Table S2**: Information about the uncirculated PacBio sequences used as reference sequences for SNP calling.

**Table S3:** Antimicrobial resistance genes and plasmid markers identified from the uncirculated PacBio sequences.

**Supplementary File 2:**

The questionnaire template in German language.

**Supplementary File 3:**

Summarized results from the questionnaire.

**Supplementary File 4:**

Sequence quality parameters for all sample sets.

**Supplementary File 5:**

CgMLST distance allele matrices for all samples.

**Supplementary File 6:**

SNP distance allele matrices for all samples.

**Supplementary File 7:** Figures show the global GC-bias across the whole genome calculated using Benjamini’s method (Benjamini and Speed, 2012) with the computeGCBias function of the deepTools package (Ramírez et al., 2016) for all sample sets. The function counts the number of reads per GC fraction and compares them to the expected GC profile, calculated by counting the number of DNA fragments per GC fraction in a reference genome. In an ideal experiment, the observed GC profile would match the expected profile, producing a flat line at 0. The fluctuations to both ends of the x-axis are due to the fact that only a small number of genome regions have extreme GC fractions.

