## Supplemental File 1 for "German-wide interlaboratory study compares consistency, accuracy and reproducibility of whole-genome short read sequencing"

**Table S1: General information about the strains used for the interlaboratory study**

| Sample | Organism | Serovar | MLST |
|---|---|---|---|
| 19-RV1-P64-1 | *Campylobacter jejuni* | | 4774 |
| 19-RV1-P64-2 | *Campylobacter jejuni* | | 21 |
| 19-RV1-P64-3 | *Listeria monocytogenes* | IIc | 9 |
| 19-RV1-P64-4 | *Listeria monocytogenes* | IIb | 59 |
| 19-RV1-P64-5 | *Salmonella enterica* subsp. *enterica* | Infantis | 32 |
| 19-RV1-P64-6 | *Salmonella enterica* subsp. *enterica* | Paratyphi B var. Java | 28 |

**Table S2: Information about the uncirculated PacBio sequences used as reference sequences for SNP calling**

| Sample | Length | Number of contigs | GC |
|---|---|---|---|
| 19-RV1-P64-1 | 1,619,699 bp | 1 | 30.53 % |
| 19-RV1-P64-2 | 1,716,034 bp | 1 | 30.51 % |
| 19-RV1-P64-3 | 3,000,034 bp | 2 | 37.96 % |
| 19-RV1-P64-4 | 3,019,943 bp | 1 | 37.93 % |
| 19-RV1-P64-5 | 5,052,870 bp | 2 | 51.98 % |
| 19-RV1-P64-6 | 4,789,411 bp | 4 | 52.22 % |

**Table S3: Antimicrobial resistance genes and plasmid markers identified from the uncirculated PacBio sequences**

| Sample | Number of plasmid markers | Plasmid markers | Number of resistance genes | Resistance genes |
|---|---|---|---|---|
| 19-RV1-P64-1 | 0 | | 2 | blaOXA-184;tet(O) |
| 19-RV1-P64-2 | 0 | | 1 | blaOXA-605 |
| 19-RV1-P64-3 | 0 | | 2 | fosX;lin |
| 19-RV1-P64-4 | 0 | | 2 | fosX;lin |
| 19-RV1-P64-5 | 3 | IncHI2_1;TrfA_1;IncHI2A_1 | 13 | sul1;qacEdelta1;aadA1;ere(A);sul1;qacEdelta1;aadA1;blaVIM-1;catA1;blaACC-1;sul1;aph(6)-Id;aph(3'')-Ib |
| 19-RV1-P64-6 | 1 | IncI1_1_Alpha | 9 | aadA1;aadA1;dfrA1;aadA1;dfrA1;sul2;blaTEM-1;dfrA1;tet(A) |
