## Supplemental File 2 for "German-wide interlaboratory study compares consistency, accuracy and reproducibility of whole-genome short read sequencing"

### Ergebnisprotokoll für den 1. §64 LFGB Ringversuch „NGS-Bakteriencharakterisierung (2019)“

durchgeführt durch das Bundesinstitut für Risikobewertung
Studienzentrum für Genomsequenzierung und –analyse
Abteilung Biologische Sicherheit

Code-Nr. des Laboratoriums :

Name des Laboratoriums :

Empfangsdatum der Ringversuchsproben :

Beginn des Ringversuches :

#### 1. Transport

##### 1.1 Wann kam das Paket mit den Ringversuchsproben an?

Datum:

##### 1.2 War das Paket beschädigt?

Ja/Nein:

#### 2. DNA Konzentrations- und Qualitätsbestimmung

##### 2.1 Welche Methode wurde für die DNA Konzentrationsbestimmung angewandt?

- Qubit
- Nanodrop
- keine
- andere:

Falls erfasst: Bitte tragen Sie die DNA Konzentration in die Excel-Ergebnisstabelle ein.

##### 2.2 Haben Sie die DNA Qualität anhand der Bestimmung von Absorptionsverhältnissen (A260/A280 und A260/A230) geprüft?

- Ja, mittels Nanodrop
- Ja, mittels:
- Nein

Falls erfasst: Bitte tragen Sie die ermittelten Werte in die Excel-Ergebnisstabelle ein.

**2.3 Wurde die Qualität (z.B. Degradation) der DNA mit Gelelektrophorese/Bioanalyzer/Fragment Analyzer überprüft?**

Ja / Nein

Falls Ja: Bitte vermerken Sie in der Excel-Ergebnisstabelle für welche DNA Proben eine Degradation beobachtet wurde.

#### 3. Bibliothekenherstellung (Library Prep)

**3.1 Welche Kits haben Sie für die Vorbereitung der Library verwendet? Bitte geben Sie den Namen und die Bestellnummer der verwendeten Kits an.**

Name:
Bestellnummer:

**3.2 Wie viel DNA haben Sie zu Beginn eingesetzt?**

Bitte tragen Sie die DNA Mengen [ng] in die Excel-Ergebnisstabelle ein.

**3.3 Erforderte Ihr Protokoll die enzymatische oder mechanische Fragmentierung der DNA?**

- enzymatisch
- mechanisch
- andere:

**3.4 In welchem Umfang folgten Sie dem Kit Protokoll?**

- vollständig
- kleine Modifikationen
  welche:
- größere Modifikationen
  welche:

**3.5 Welche Methode wurde zum Überprüfen der Qualität/Fragmentlänge der erzeugten Bibliothek verwendet?**

- Bioanalyzer/ Fragment Analyzer

- Gelelektrophorese
- keine
- andere:

**3.6 Haben Sie die DNA Bibliotheken normalisiert?**

Ja / Nein

Falls ja, über welche Methode:

- manuell über die Konzentration und Fragmentlänge
  mit Konzentrationsbestimmung über:
    - Qubit
    - qPCR
- Bead-basiert
- andere:

**3.7 Wie wurde die DNA gepooled?**

- alle Proben gleich
- gewichtet gemäß der Genomgröße der unterschiedlichen Spezies
- andere:

**3.8 Was war die Ausgangsgesamtgröße aller sequenzierten Genome in Millionen Basenpaaren (Mb) in ihrem Sequenzierungslauf*?**

Gesamtgröße [Mb]:

* Hinweis: 1 *Salmonella* Genom: ca. 5 Mb
1 *Campylobacter* Genom: ca. 1,7 Mb
1 *Listeria* Genom: ca. 3 Mb
2 x *Salmonella*, 2 x *Listeria*, 2 x *Campylobacter*: 19,4 Mb
24 x Salmonella: 120 Mb
etc…

**3.9 Welche Library-Menge haben Sie für die Sequenzierung eingesetzt?**

Illumina- manuelle Normalisierung:
- Library-Ladekonzentration [pM]:

Illumina- bead-basierte Normalisierung
- Volumen des Library Pools [µl]:

Ion Torrent (Ladekonzentration für emPCR)
- Library-Ladekonzentration [pM]:
- weitere relevante Angaben:

#### 4. Sequenzierung

##### 4.1 Welche Sequenzierungsplattform nutzen Sie?

Illumina
- iSeq
- MiniSeq
- MiSeq series
- NextSeq series
- HiSeq series
- NovaSeq series

Ion Torrent
- PGM
- Proton
- S5/S5XL

andere:

##### 4.2 Verwenden Sie eine Sequenzierungskontrolle?
(z.B. PhiX, Test Fragments, Standard-DNA, definierte Isolat-DNA, etc.)

Ja/Nein

Falls ja, welche:

##### 4.3 Welches Kit verwenden Sie für die Sequenzierung der Proben? Bitte geben Sie den Namen und die Bestellnummer des verwendeten Kits an.

Name:
Bestellnummer:

##### 4.4 Welche Sequenzierungstiefe (x-fache Abdeckung der Genome) streben Sie in der Regel an?

Sequenzierungstiefe:

##### 4.5 Weitere Fragen für die Sequenzierung mittels Illumina Geräten

###### 4.5.1 Sequenzieren Sie single-end oder paired-end?

- single-end
- paired-end

###### 4.5.2 Wie viele Zyklen sequenzieren Sie?
(Für die paired-end Sequenzierung bitte mit 2x angeben, z.B. 2x151 Zyklen)

Zyklenzahl:

##### 4.5.3 Welche Cluster-Dichte haben Sie erreicht?

K/mm2:

##### 4.5.4 Wie groß war die Prozentzahl an Basen mit einem Qualitäts-Score >=30?

>=Q30%:

##### 4.5.5 Wie hoch war die Gesamtausbeute an sequenzierten Basen (total yield [G])?

Total yield [G]:

##### 4.5.6 Nennung weiterer relevanter Qualitätsparameter, die Sie zur Qualitätsprüfung eines Laufes heranziehen:

#### 4.6 Weitere Fragen für die Sequenzierung mittels Ion Torrent Geräten

##### 4.6.1 Wie hoch war ihre ISP Density und wie viele ISPs waren polyklonal?

ISP Density:
polyklonal [%]:

##### 4.6.2 Wie groß ist die totale Readanzahl (total reads), der Prozentsatz an verwendbaren Reads und die durchschnittliche Readlänge?

Total reads:
usable reads (%):
durchschnittliche Readlänge:

##### 4.6.3 Nennung weiterer relevanter Qualitätsparameter, die Sie zur Qualitätsprüfung eines Laufes heranziehen:

#### 5. Anmerkungen zur Ringversuchsdurchführung

Anmerkungen zu Faktoren und Problemen, die das Ergebnis beeinflusst haben könnten:

#### 6. Anmerkungen zur Ringversuchsorganisation und Verbesserungsvorschläge

- Probenversand:
- Ergebnisdokumentation:
- Datenübermittlung:
- Sonstiges:
