## Supplemental File 7 for "German-wide interlaboratory study compares consistency, accuracy and reproducibility of whole-genome short read sequencing"

# LC02a

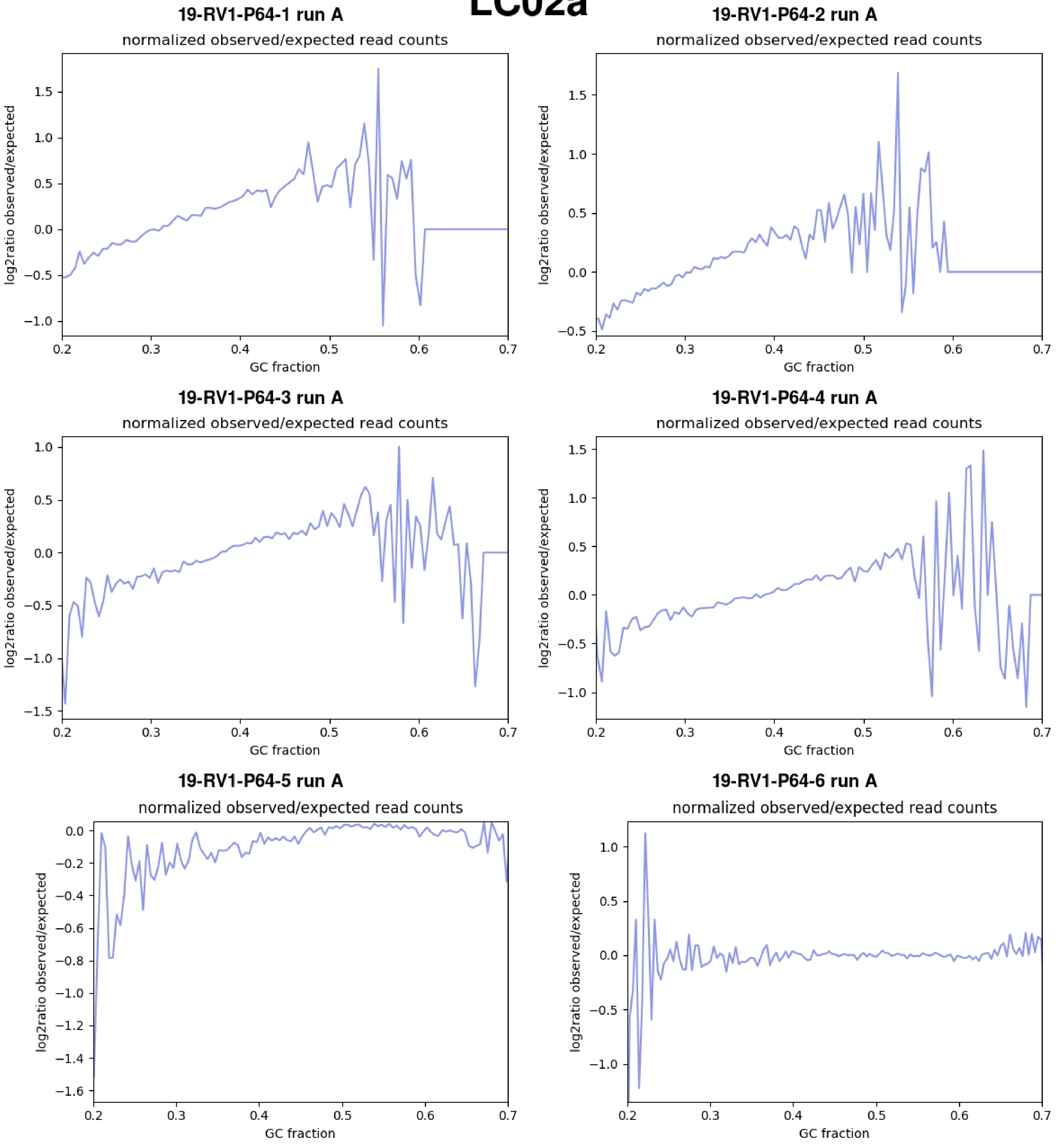
19-RV1-P64-1 run A
normalized observed/expected read counts
log2ratio observed/expected
GC fraction
19-RV1-P64-2 run A
normalized observed/expected read counts
log2ratio observed/expected
GC fraction
19-RV1-P64-3 run A
normalized observed/expected read counts
log2ratio observed/expected
GC fraction
19-RV1-P64-4 run A
normalized observed/expected read counts
log2ratio observed/expected
GC fraction
19-RV1-P64-5 run A
normalized observed/expected read counts
log2ratio observed/expected
GC fraction
19-RV1-P64-6 run A
normalized observed/expected read counts
log2ratio observed/expected
GC fraction

# LC02b

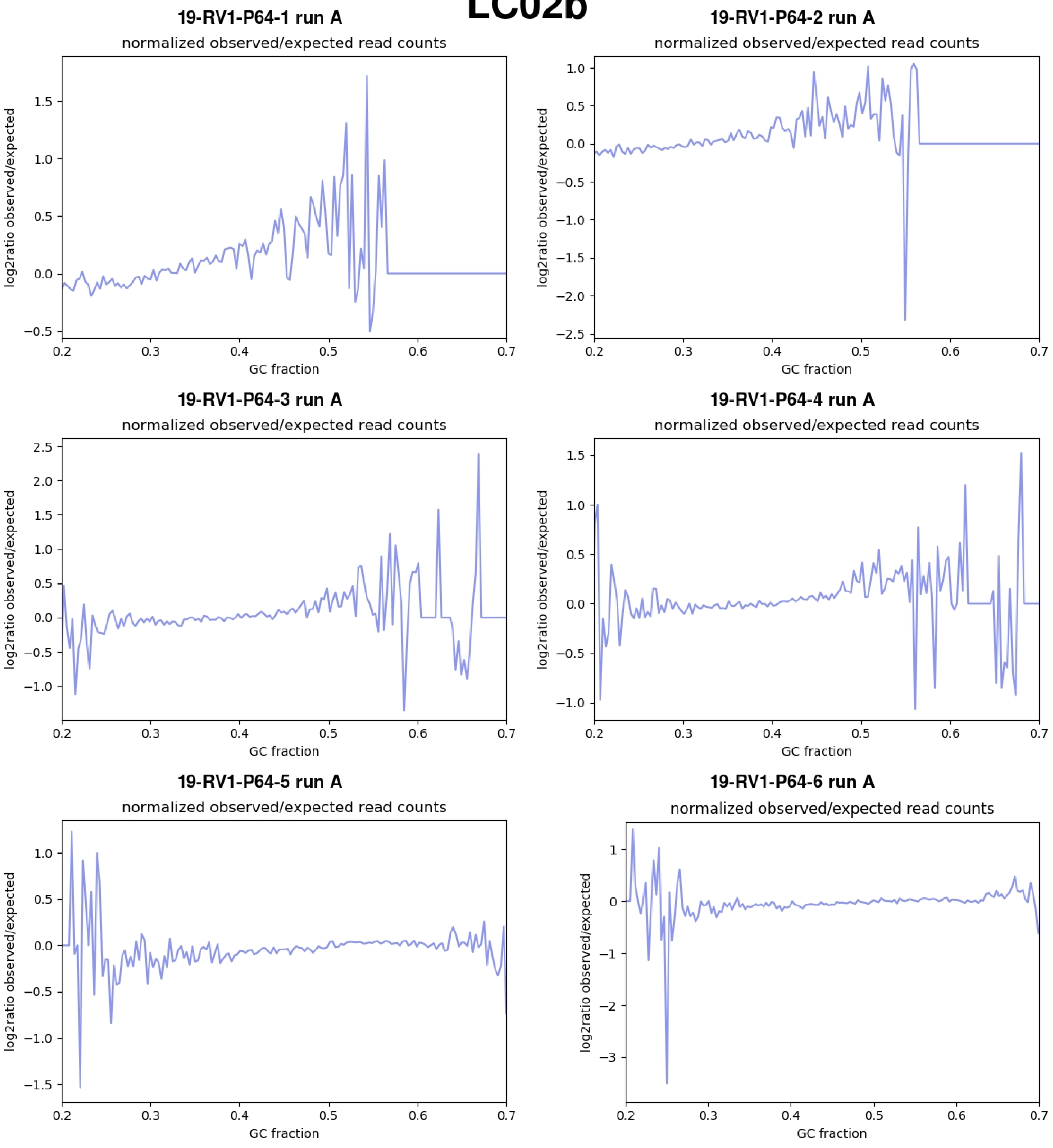
19-RV1-P64-1 run A
normalized observed/expected read counts
19-RV1-P64-2 run A
normalized observed/expected read counts
19-RV1-P64-3 run A
normalized observed/expected read counts
19-RV1-P64-4 run A
normalized observed/expected read counts
19-RV1-P64-5 run A
normalized observed/expected read counts
19-RV1-P64-6 run A
normalized observed/expected read counts
log2ratio observed/expected
GC fraction

# LC02c

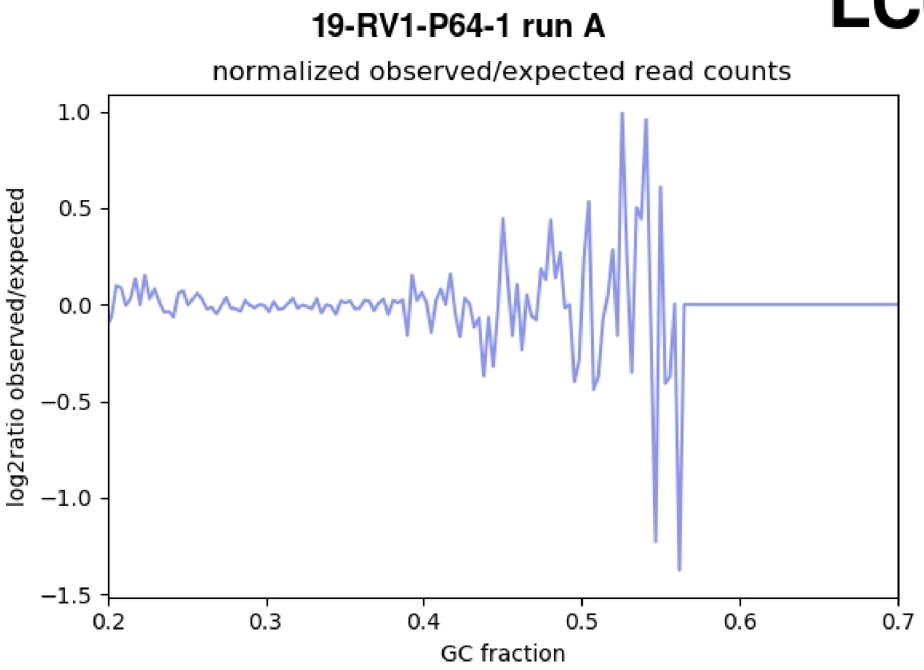
19-RV1-P64-1 run A
normalized observed/expected read counts
log2ratio observed/expected
1.0
0.5
0.0
−0.5
−1.0
−1.5
0.2
0.3
0.4
0.5
0.6
0.7
GC fraction

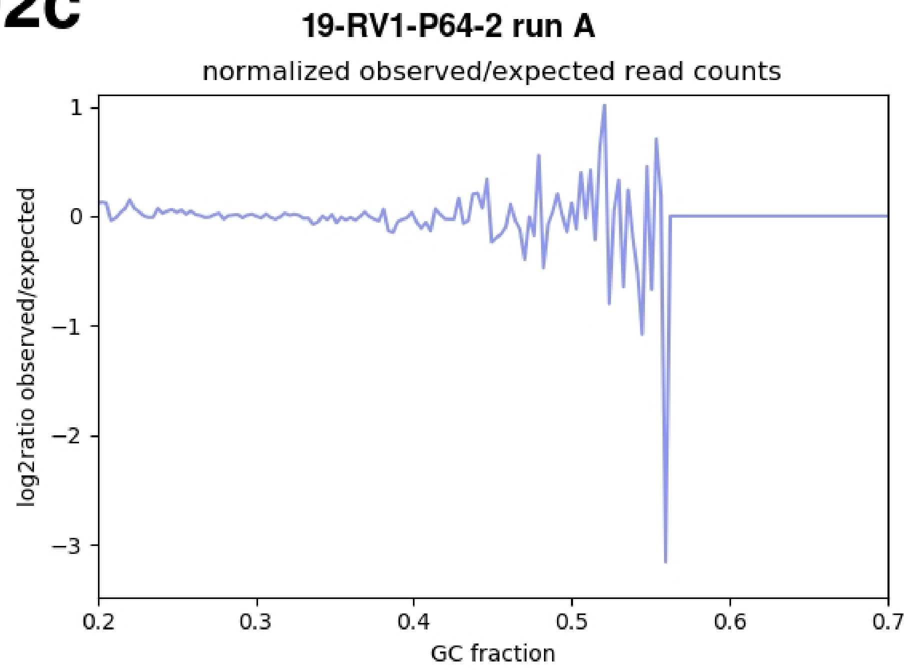
19-RV1-P64-2 run A
normalized observed/expected read counts
log2ratio observed/expected
1
0
−1
−2
−3
0.2
0.3
0.4
0.5
0.6
0.7
GC fraction

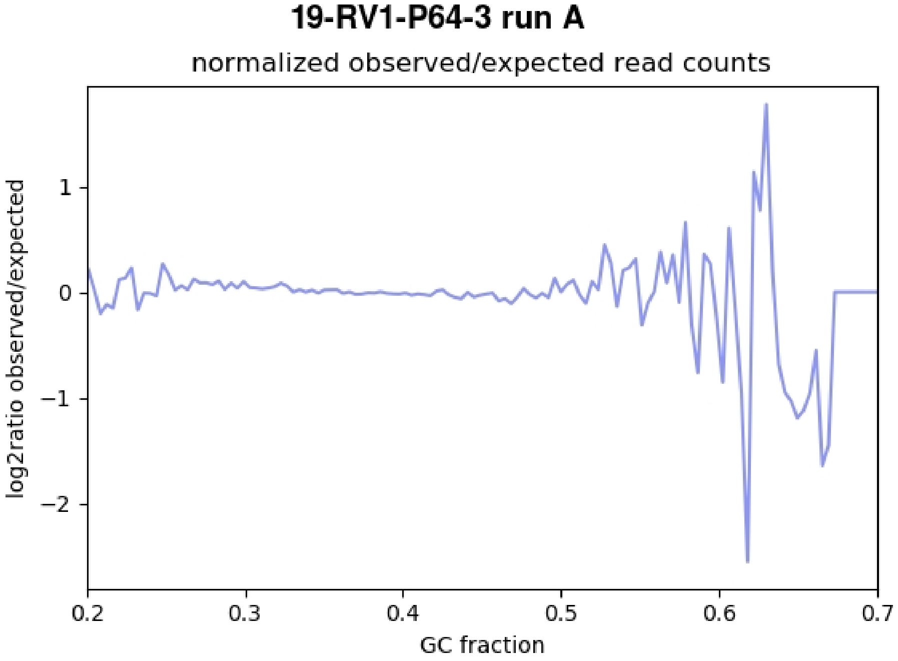
19-RV1-P64-3 run A
normalized observed/expected read counts
log2ratio observed/expected
1
0
−1
−2
0.2
0.3
0.4
0.5
0.6
0.7
GC fraction

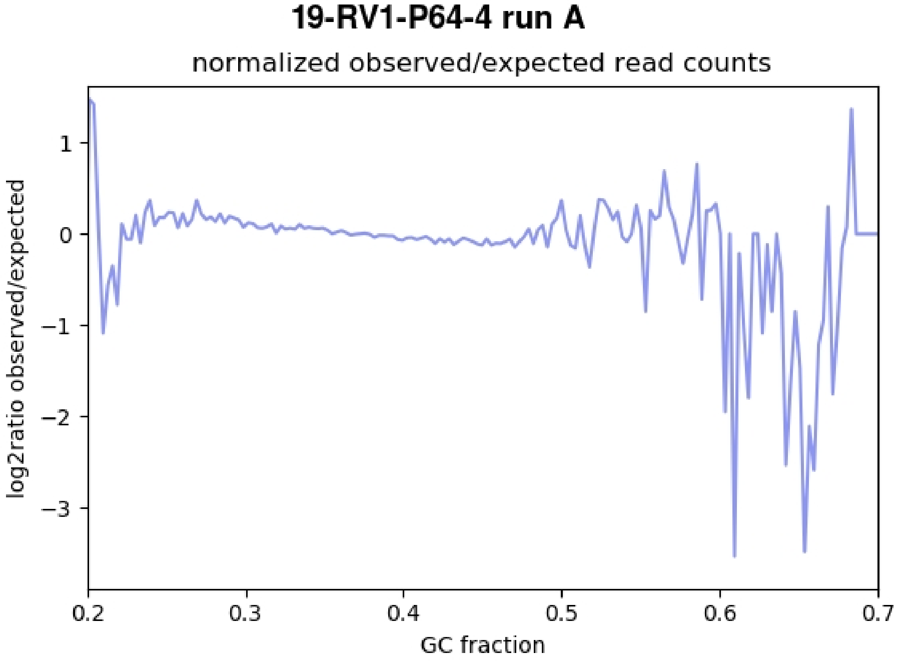
19-RV1-P64-4 run A
normalized observed/expected read counts
log2ratio observed/expected
1
0
−1
−2
−3
0.2
0.3
0.4
0.5
0.6
0.7
GC fraction

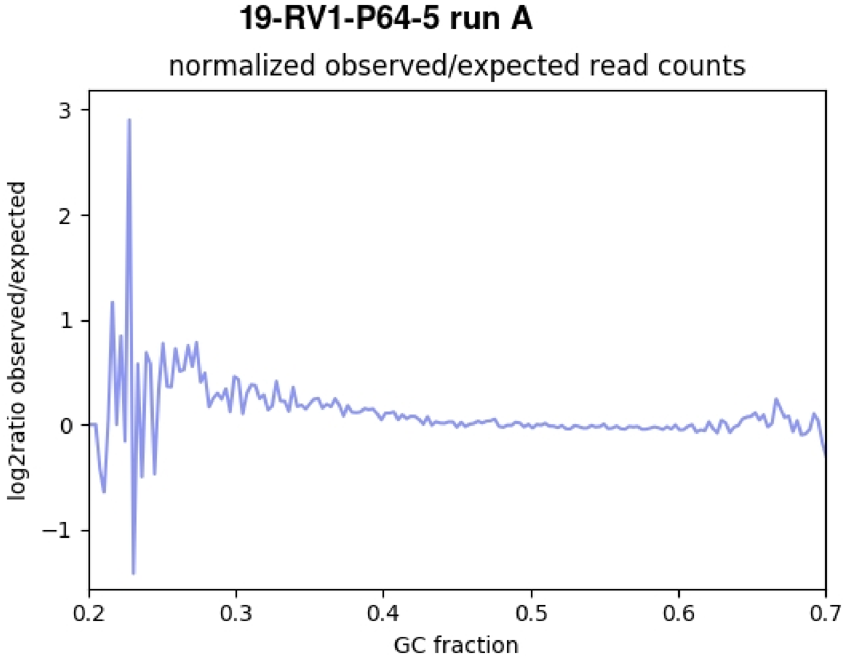
19-RV1-P64-5 run A
normalized observed/expected read counts
log2ratio observed/expected
3
2
1
0
−1
0.2
0.3
0.4
0.5
0.6
0.7
GC fraction

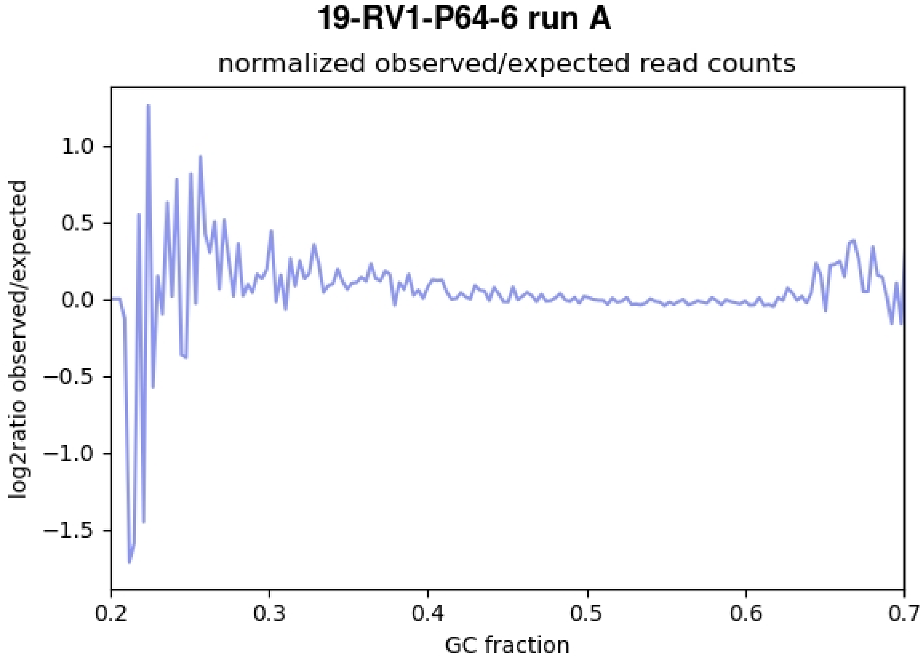
19-RV1-P64-6 run A
normalized observed/expected read counts
log2ratio observed/expected
1.0
0.5
0.0
−0.5
−1.0
−1.5
0.2
0.3
0.4
0.5
0.6
0.7
GC fraction

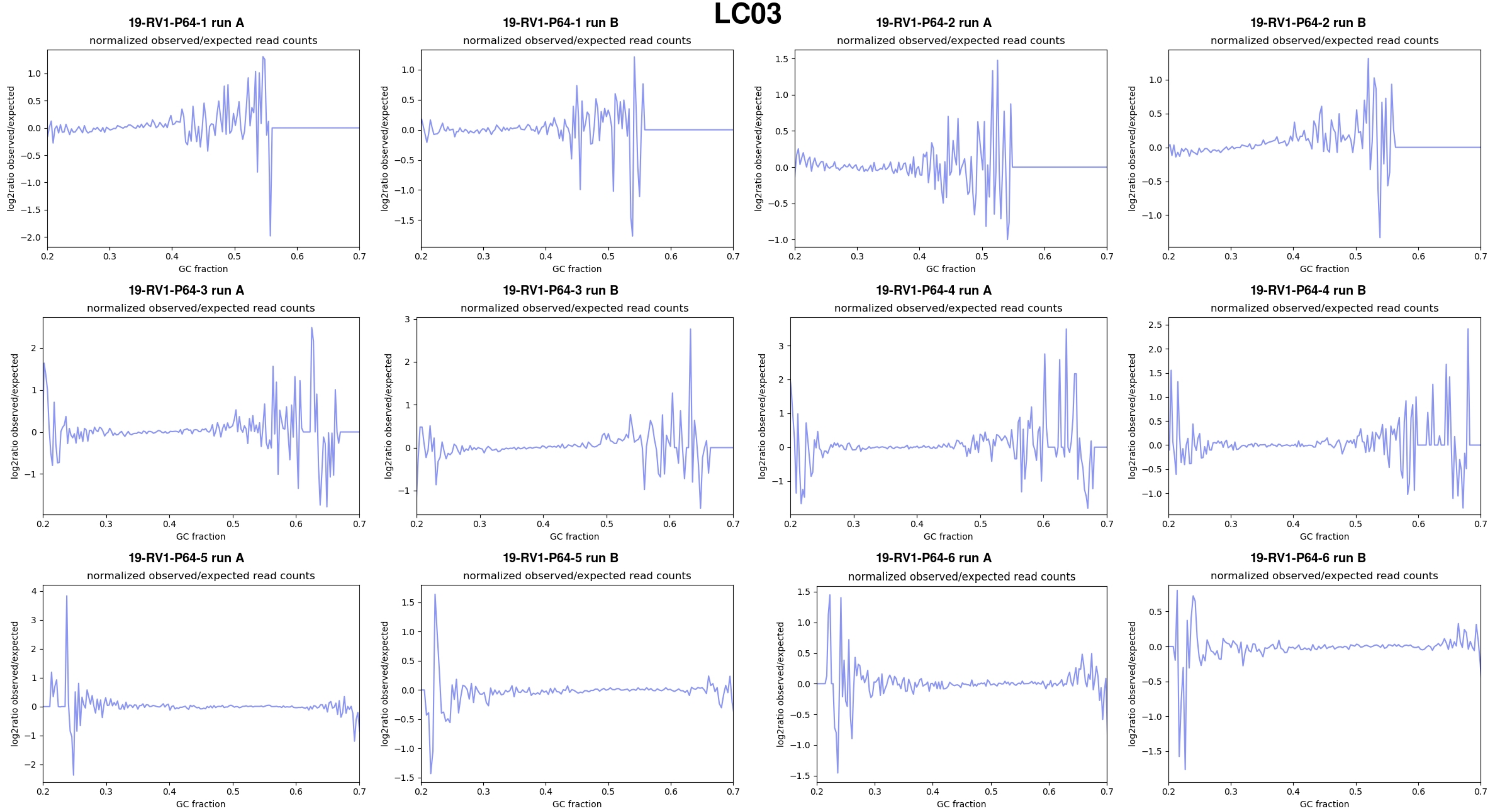
LC03
19-RV1-P64-1 run A
normalized observed/expected read counts
19-RV1-P64-1 run B
normalized observed/expected read counts
19-RV1-P64-2 run A
normalized observed/expected read counts
19-RV1-P64-2 run B
normalized observed/expected read counts
19-RV1-P64-3 run A
normalized observed/expected read counts
19-RV1-P64-3 run B
normalized observed/expected read counts
19-RV1-P64-4 run A
normalized observed/expected read counts
19-RV1-P64-4 run B
normalized observed/expected read counts
19-RV1-P64-5 run A
normalized observed/expected read counts
19-RV1-P64-5 run B
normalized observed/expected read counts
19-RV1-P64-6 run A
normalized observed/expected read counts
19-RV1-P64-6 run B
normalized observed/expected read counts
log2ratio observed/expected
GC fraction

# LC04

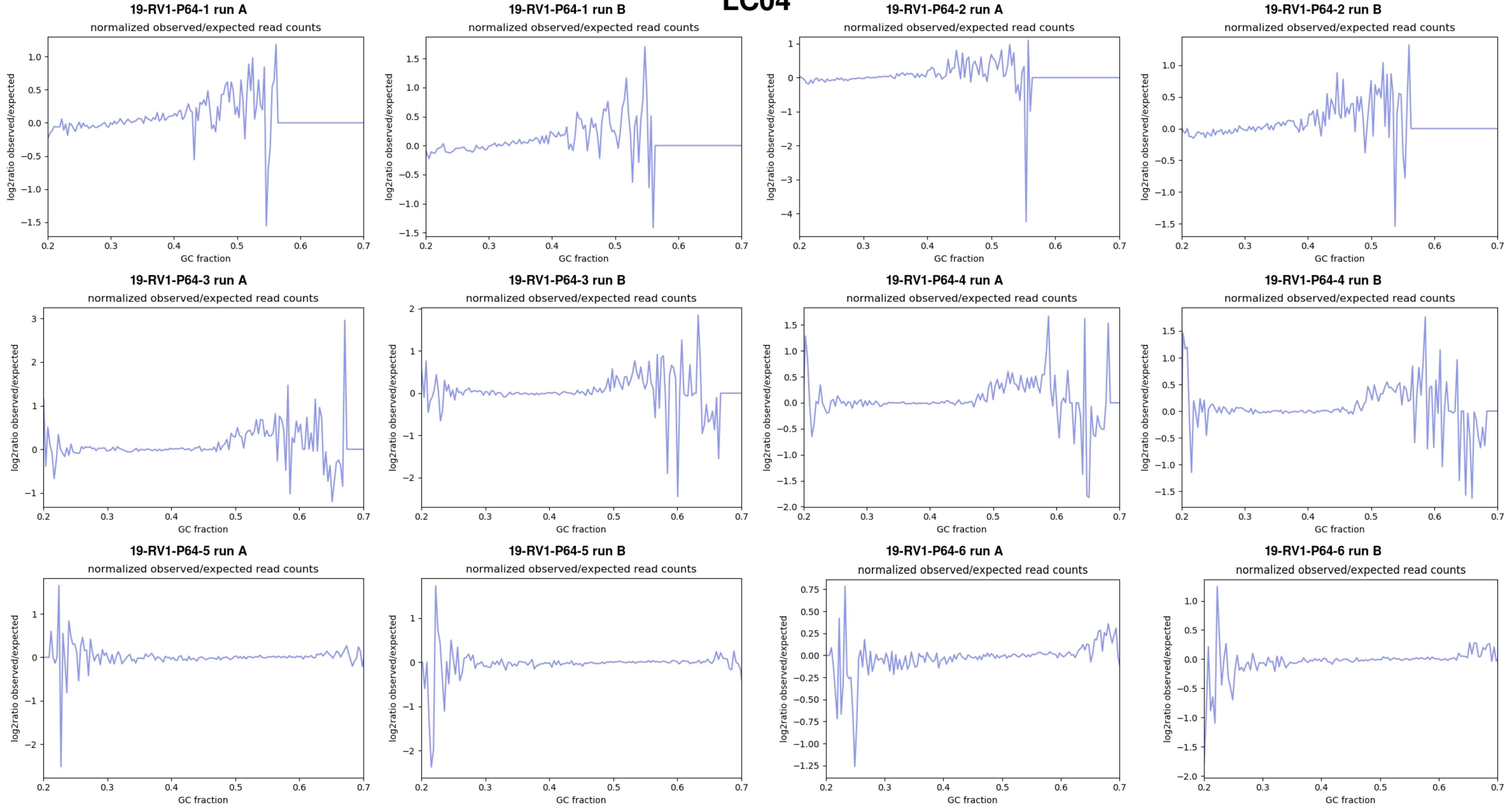
19-RV1-P64-1 run A
normalized observed/expected read counts
19-RV1-P64-1 run B
normalized observed/expected read counts
19-RV1-P64-2 run A
normalized observed/expected read counts
19-RV1-P64-2 run B
normalized observed/expected read counts
19-RV1-P64-3 run A
normalized observed/expected read counts
19-RV1-P64-3 run B
normalized observed/expected read counts
19-RV1-P64-4 run A
normalized observed/expected read counts
19-RV1-P64-4 run B
normalized observed/expected read counts
19-RV1-P64-5 run A
normalized observed/expected read counts
19-RV1-P64-5 run B
normalized observed/expected read counts
19-RV1-P64-6 run A
normalized observed/expected read counts
19-RV1-P64-6 run B
normalized observed/expected read counts
log2ratio observed/expected
GC fraction

# LC05

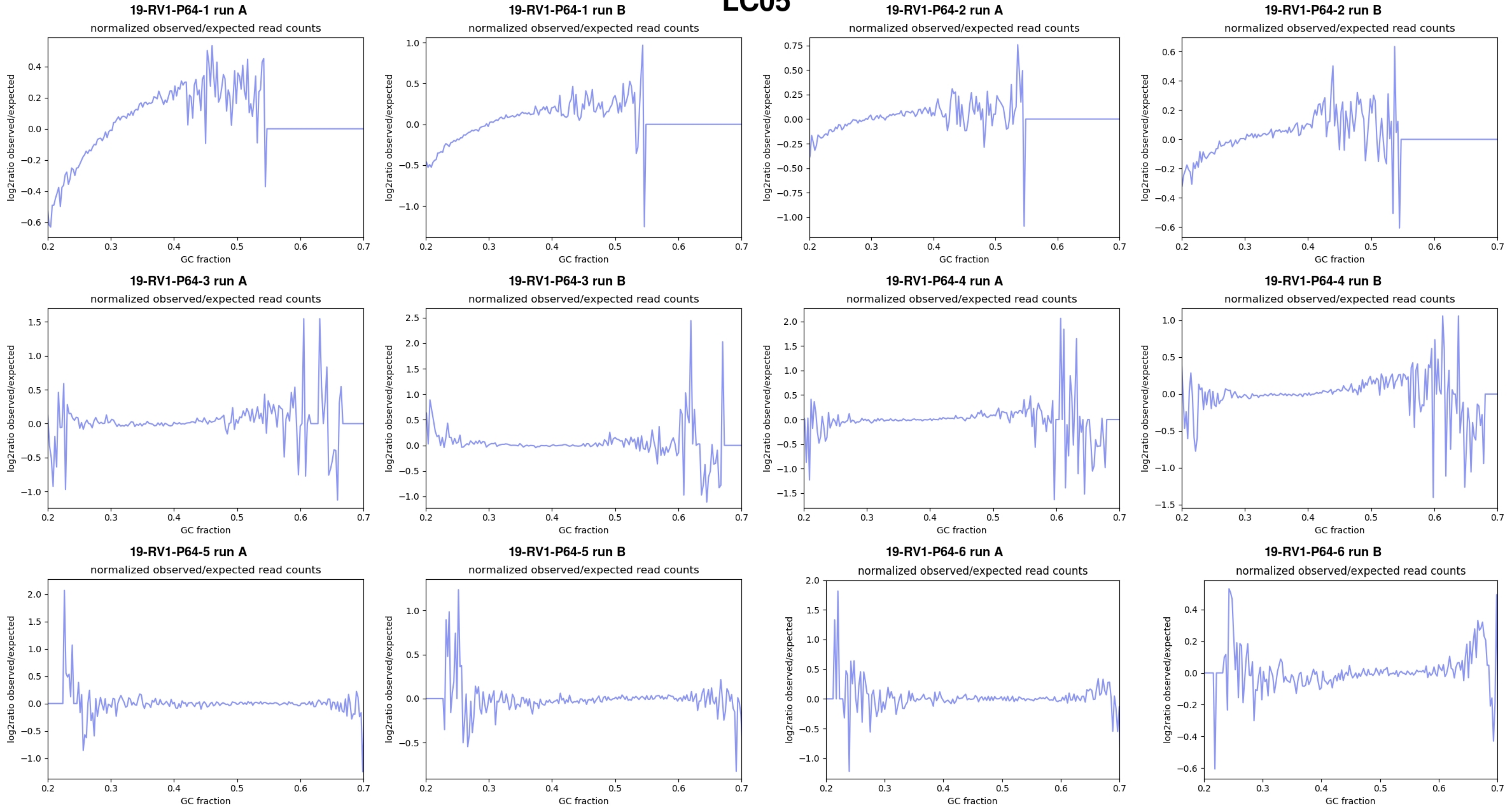
19-RV1-P64-1 run A
normalized observed/expected read counts
19-RV1-P64-1 run B
normalized observed/expected read counts
19-RV1-P64-2 run A
normalized observed/expected read counts
19-RV1-P64-2 run B
normalized observed/expected read counts
19-RV1-P64-3 run A
normalized observed/expected read counts
19-RV1-P64-3 run B
normalized observed/expected read counts
19-RV1-P64-4 run A
normalized observed/expected read counts
19-RV1-P64-4 run B
normalized observed/expected read counts
19-RV1-P64-5 run A
normalized observed/expected read counts
19-RV1-P64-5 run B
normalized observed/expected read counts
19-RV1-P64-6 run A
normalized observed/expected read counts
19-RV1-P64-6 run B
normalized observed/expected read counts
log2ratio observed/expected
GC fraction

# LC06

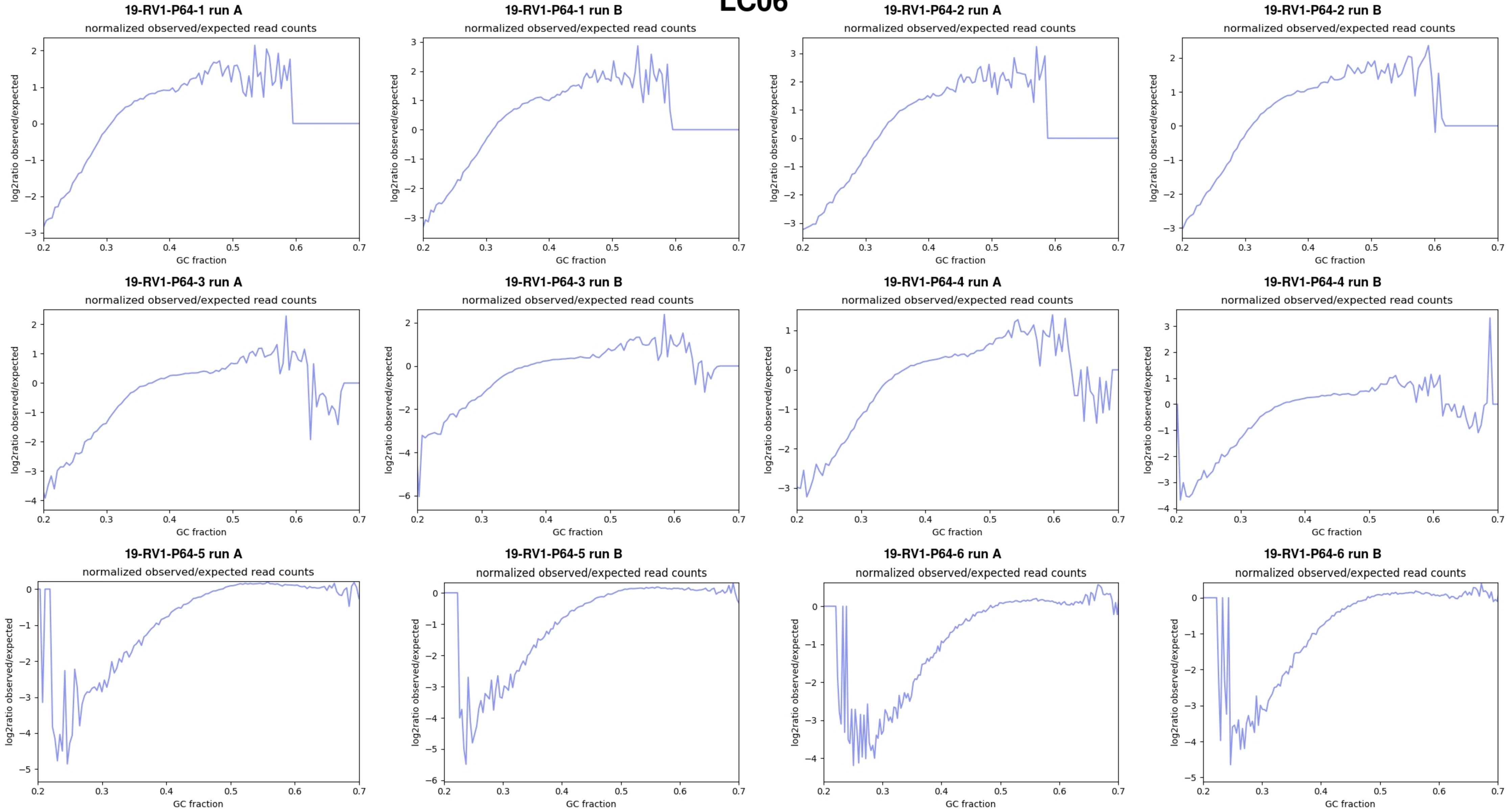
19-RV1-P64-1 run A
normalized observed/expected read counts
log2ratio observed/expected
GC fraction
19-RV1-P64-1 run B
normalized observed/expected read counts
log2ratio observed/expected
GC fraction
19-RV1-P64-2 run A
normalized observed/expected read counts
log2ratio observed/expected
GC fraction
19-RV1-P64-2 run B
normalized observed/expected read counts
log2ratio observed/expected
GC fraction
19-RV1-P64-3 run A
normalized observed/expected read counts
log2ratio observed/expected
GC fraction
19-RV1-P64-3 run B
normalized observed/expected read counts
log2ratio observed/expected
GC fraction
19-RV1-P64-4 run A
normalized observed/expected read counts
log2ratio observed/expected
GC fraction
19-RV1-P64-4 run B
normalized observed/expected read counts
log2ratio observed/expected
GC fraction
19-RV1-P64-5 run A
normalized observed/expected read counts
log2ratio observed/expected
GC fraction
19-RV1-P64-5 run B
normalized observed/expected read counts
log2ratio observed/expected
GC fraction
19-RV1-P64-6 run A
normalized observed/expected read counts
log2ratio observed/expected
GC fraction
19-RV1-P64-6 run B
normalized observed/expected read counts
log2ratio observed/expected
GC fraction

# LC07

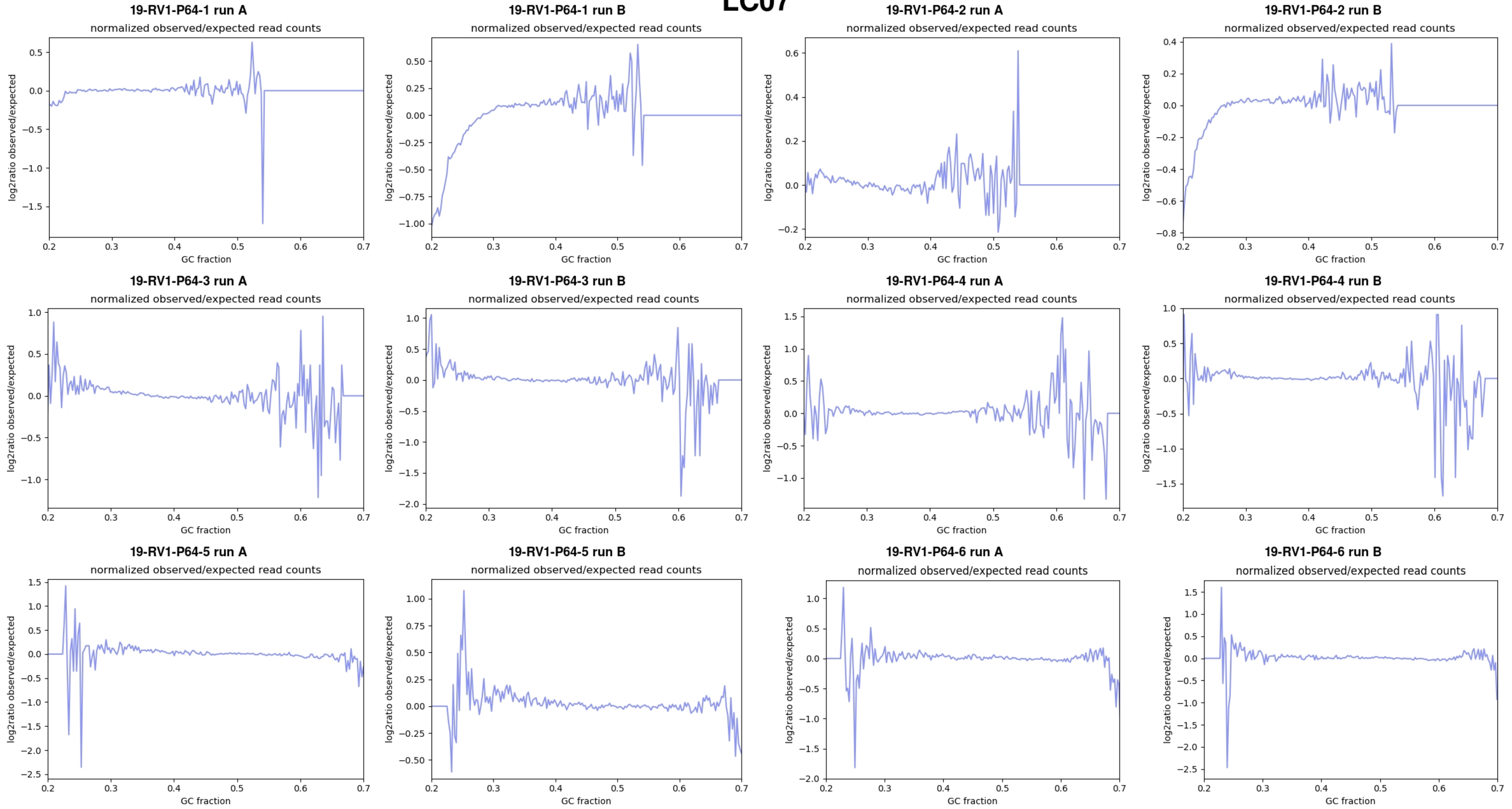
19-RV1-P64-1 run A
19-RV1-P64-1 run B
19-RV1-P64-2 run A
19-RV1-P64-2 run B
19-RV1-P64-3 run A
19-RV1-P64-3 run B
19-RV1-P64-4 run A
19-RV1-P64-4 run B
19-RV1-P64-5 run A
19-RV1-P64-5 run B
19-RV1-P64-6 run A
19-RV1-P64-6 run B
normalized observed/expected read counts
log2ratio observed/expected
GC fraction

# LC08

**19-RV1-P64-1 run A**

normalized observed/expected read counts

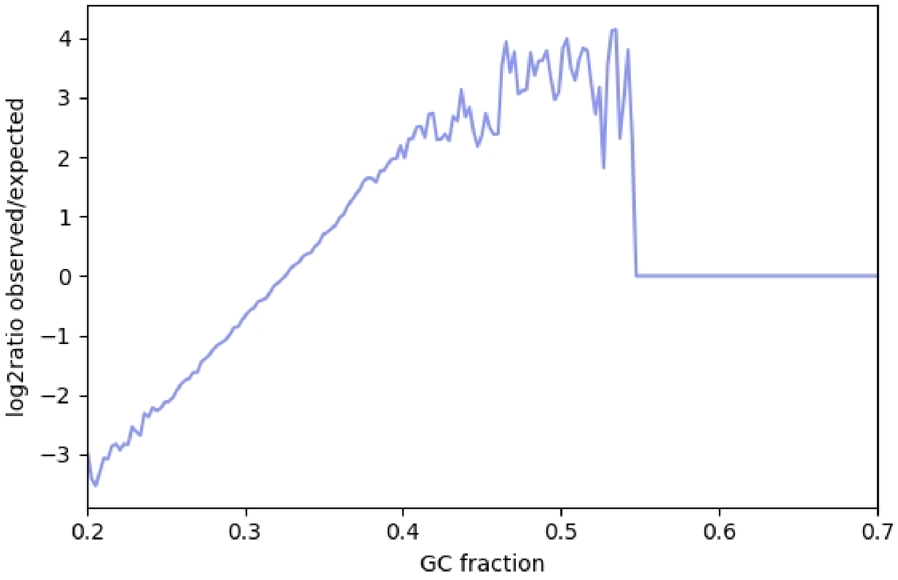


**19-RV1-P64-2 run A**

normalized observed/expected read counts

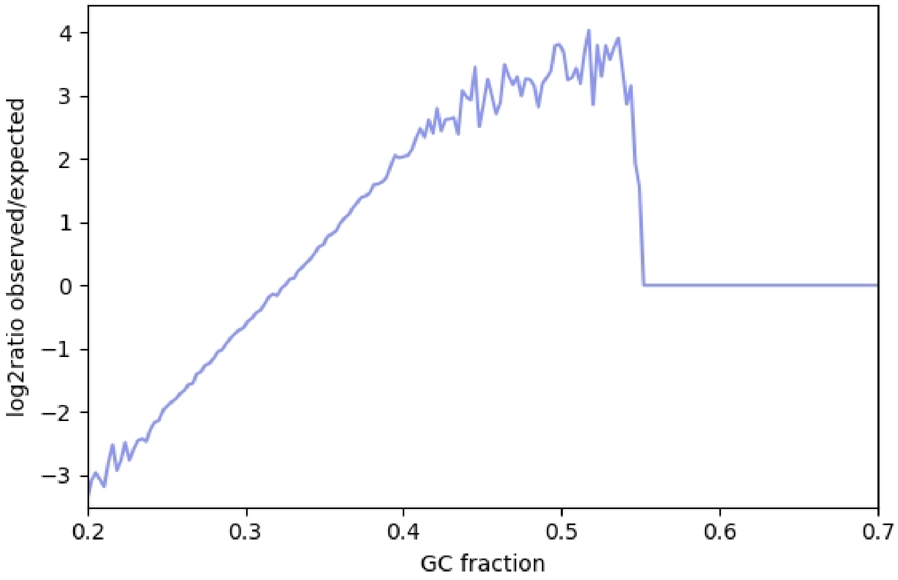


**19-RV1-P64-3 run A**

normalized observed/expected read counts

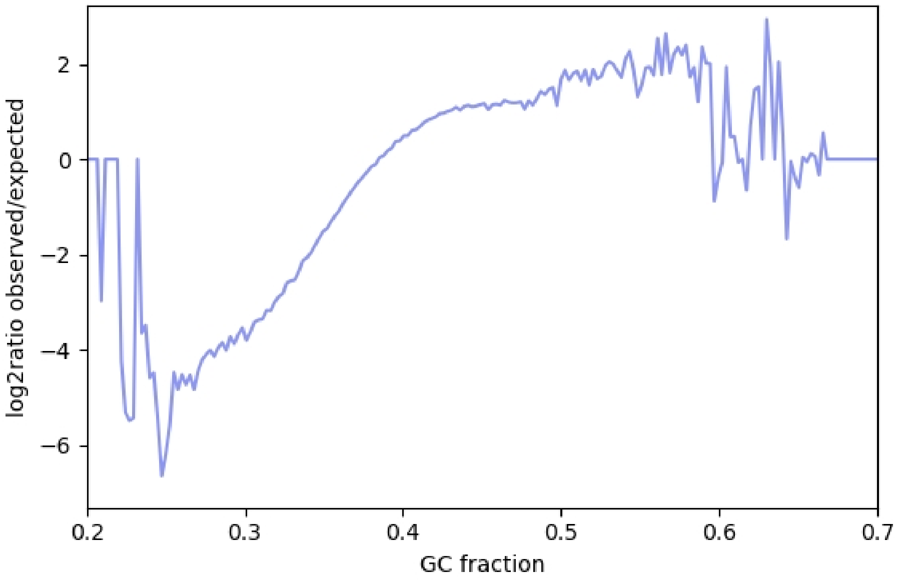


**19-RV1-P64-4 run A**

normalized observed/expected read counts

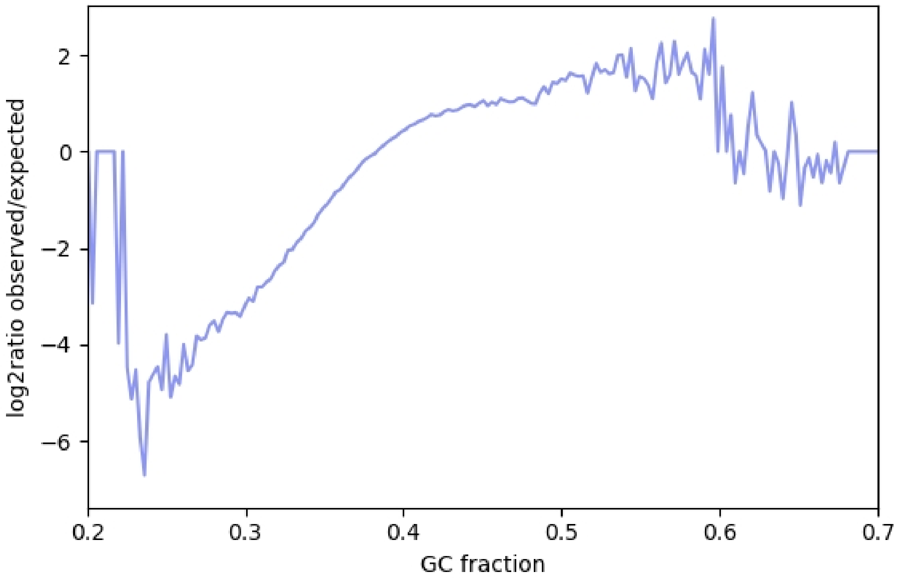


**19-RV1-P64-5 run A**

normalized observed/expected read counts

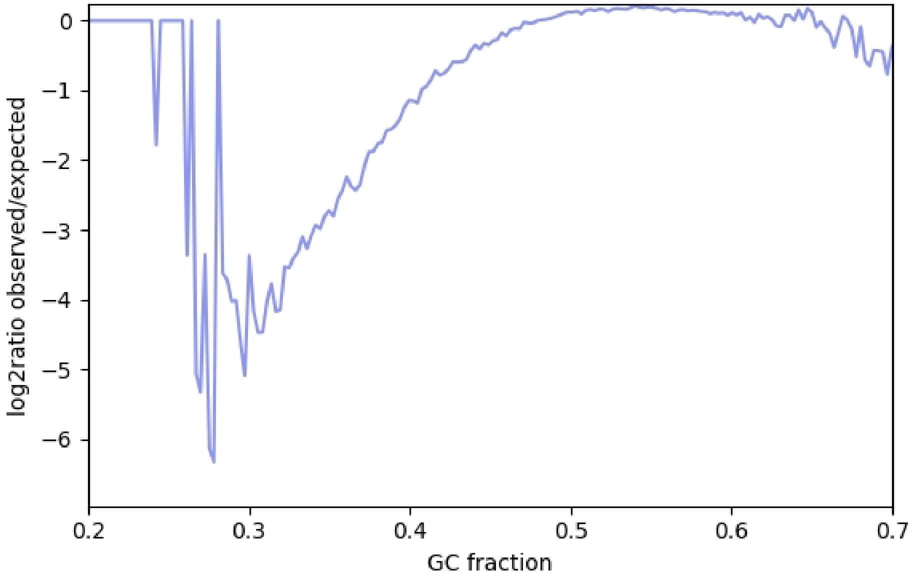


**19-RV1-P64-6 run A**

normalized observed/expected read counts

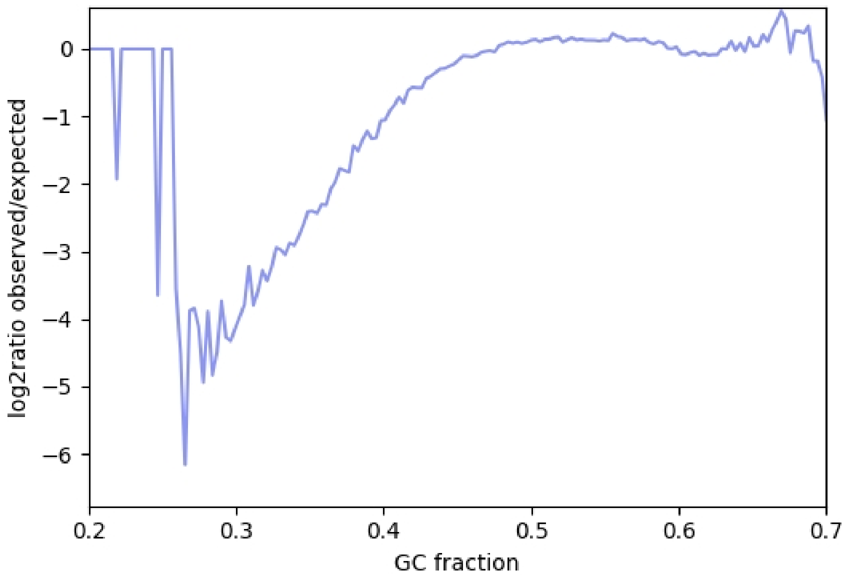

# LC09

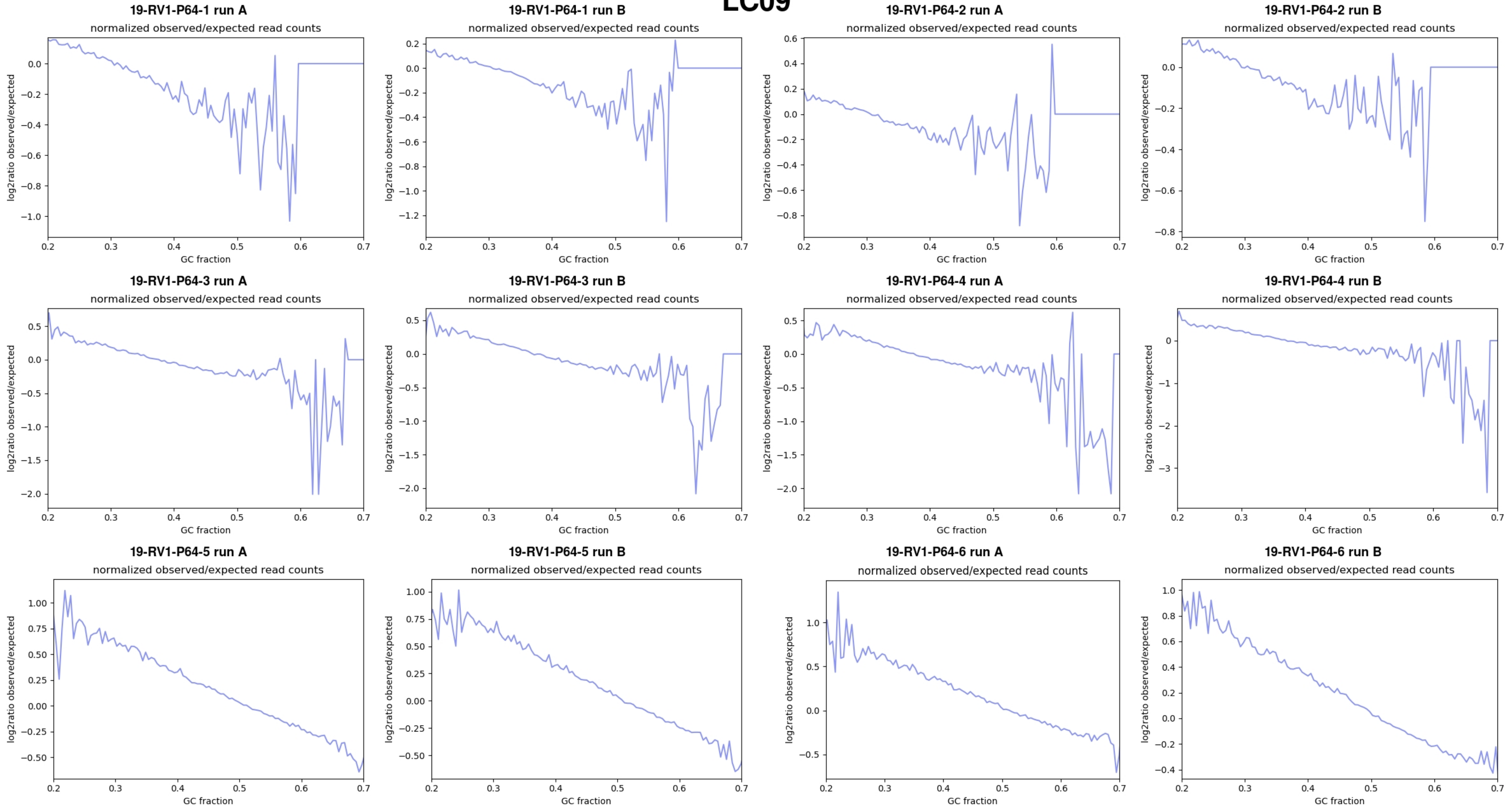
19-RV1-P64-1 run A
normalized observed/expected read counts
19-RV1-P64-1 run B
normalized observed/expected read counts
19-RV1-P64-2 run A
normalized observed/expected read counts
19-RV1-P64-2 run B
normalized observed/expected read counts
19-RV1-P64-3 run A
normalized observed/expected read counts
19-RV1-P64-3 run B
normalized observed/expected read counts
19-RV1-P64-4 run A
normalized observed/expected read counts
19-RV1-P64-4 run B
normalized observed/expected read counts
19-RV1-P64-5 run A
normalized observed/expected read counts
19-RV1-P64-5 run B
normalized observed/expected read counts
19-RV1-P64-6 run A
normalized observed/expected read counts
19-RV1-P64-6 run B
normalized observed/expected read counts
log2ratio observed/expected
GC fraction

# LC10

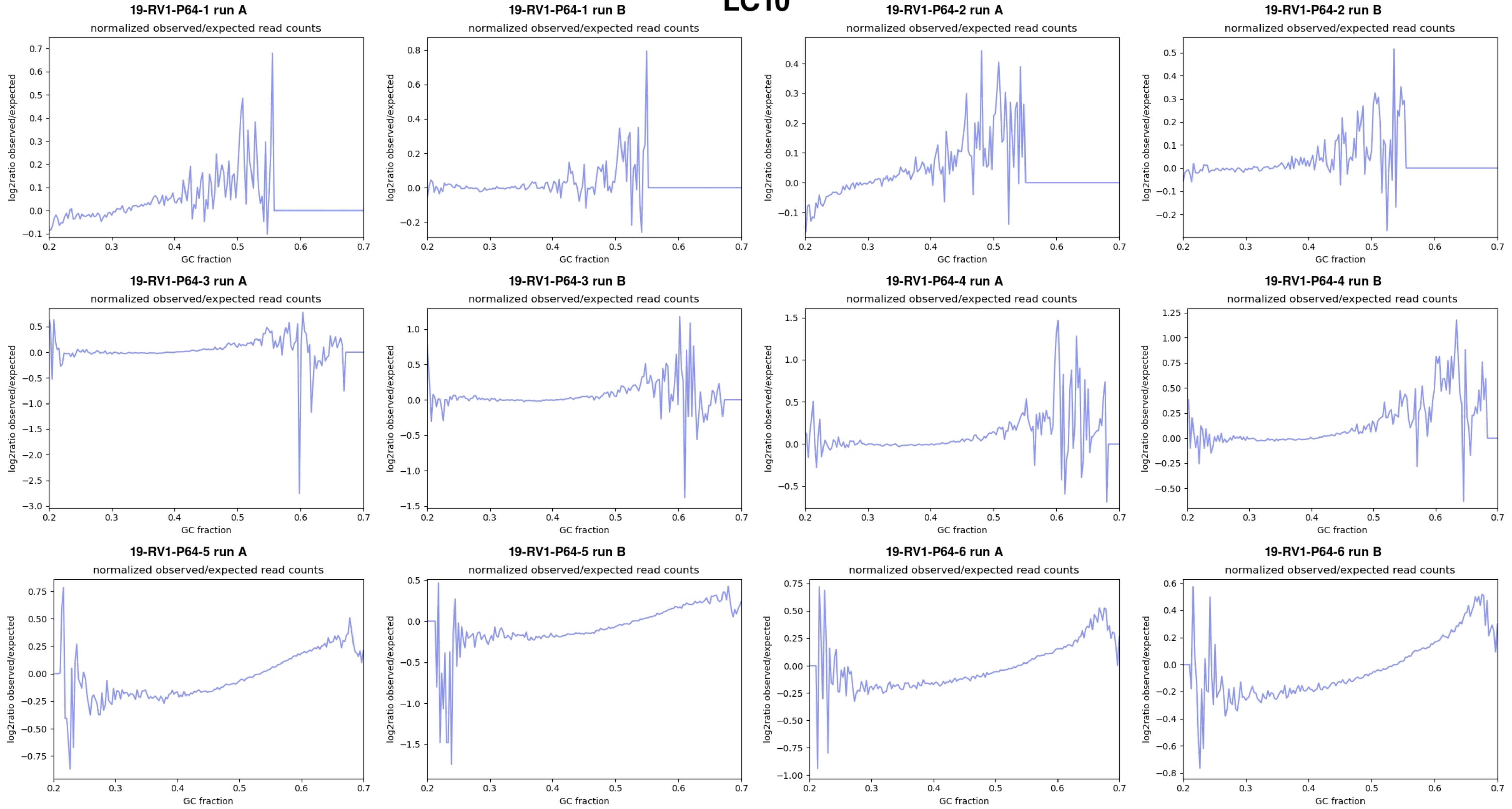

19-RV1-P64-1 run A
normalized observed/expected read counts
19-RV1-P64-1 run B
normalized observed/expected read counts
19-RV1-P64-2 run A
normalized observed/expected read counts
19-RV1-P64-2 run B
normalized observed/expected read counts
19-RV1-P64-3 run A
normalized observed/expected read counts
19-RV1-P64-3 run B
normalized observed/expected read counts
19-RV1-P64-4 run A
normalized observed/expected read counts
19-RV1-P64-4 run B
normalized observed/expected read counts
19-RV1-P64-5 run A
normalized observed/expected read counts
19-RV1-P64-5 run B
normalized observed/expected read counts
19-RV1-P64-6 run A
normalized observed/expected read counts
19-RV1-P64-6 run B
normalized observed/expected read counts
log2ratio observed/expected
GC fraction
